# A Chromodomain Mutation Identifies Separable Roles for *C. elegans* MRG-1 in Germline and Somatic Development

**DOI:** 10.1101/2022.02.19.479917

**Authors:** Christine A. Doronio, Huiping Ling, Elizabeth J. Gleason, William G. Kelly

## Abstract

The packaging of DNA into chromatin strongly influences gene regulation. Post-translational modifications of histones, and the proteins that bind to them, alter the accessibility of chromatin and contribute to the activation and repression of genes. The human MRG15 (*MORF4-* Related Gene on chromosome 15) protein is a conserved chromodomain-containing protein that binds to methylated lysine 36 on histone H3 (H3K36me) and plays important roles in development, genome integrity, and gene regulation. MRG15 affects transcriptional regulation through its interactions with both histone acetyltransferase (HAT) and histone deacetylase (HDAC) complexes. MRG-1, its *C. elegans* homolog, has similarly been shown to have important roles in genomic integrity and development, and has also been shown to co-purify with HDAC complexes. Like MRG15, MRG-1 is predicted to bind to H3K36me through its chromodomain, yet despite *mrg-1* mutants displaying developmental and germline phenotypes that overlap with H3K36 methyltransferase mutants, the role of the MRG-1 chromodomain has never been characterized. In this study, we examined meiotic cells lacking H3K36me3 to compare to *mrg-1* mutant germ cell phenotypes, and mutated key residues in the MRG-1 chromodomain (CD) to assess its function. The CD mutations cause embryonic lethality but few post-embryonic germline defects, in contrast to *mrg-1* deletion mutants which are viable but sterile. The CD mutations therefore disrupt somatic development despite the apparent absence of a requirement for MRG-1 protein in embryogenesis. Furthermore, the CD mutants exhibit a dominant RNAi resistance phenotype that is not seen in other *mrg-1* mutant alleles. This suggests that the function of MRG-1, and the chromatin modifying complexes with which it interacts, includes tissue-specific interactions involving different requirements for a functional chromodomain. We propose that the CD mutation disrupts proper guidance of complexes within which it acts, and this guidance defect results in improper HDAC and/or HAT regulation causing an indirect defect in RNAi machinery expression or targeting.

## Introduction

The regulation of genes is largely controlled by their structural accessibility to transcription machinery within the nucleus. The structural consequences of the architecture within which DNA is packaged by nucleosomes in chromatin contributes to gene activation and repression, with different epigenetic modifications such as histone modifications and DNA methylation affecting that architecture. Consequently, proteins that alter and interact with chromatin play a major role in proper gene regulation and organismal development.

In humans, MRG15 (*MORF*4- Related Gene on Chromosome 15) interacts with the histone modification tri-methylated lysine 36 on Histone H3 (H3K36me3) through MRG15’s conserved chromodomain. Chromatin binding proteins like MRG15 that recognize specific histone modifications are referred to as epigenetic “readers” and are often associated with larger chromatin remodeling complexes that can alter transcription states of genes. For example, MRG15 is essential for the assembly of MAF1 and MAF2, HAT related complexes necessary for the activation of the B-myb promoter (1). Additionally, MRG15 is required for histone acetylation and subsequent RNA Polymerase II recruitment to lipid metabolism genes in mice (2). The presence of MRG15 at certain promoters can also prevent the binding of HDAC complexes during the cell cycle (3). Interestingly, MRG15 has been shown to play a role in both histone acetyltransferase (HAT) and histone deacetylase (HDAC) complexes that are associated with gene activation and repression respectively (3, 4). The requirement for MRG15 in various chromatin remodeling complexes suggests that its ability to “read” epigenetic modifications influences chromatin-regulated differentiation in specific cell types during embryonic development in mammals (4, 5).

Perhaps related to its role in chromatin binding and remodeling, MRG15 is known to contribute to DNA repair and genomic integrity. Through its interaction with the BRCA complex, MRG15 recruits different DNA damage response proteins, like RAD51 and PALB2, during homology directed DNA repair (6). Similar roles for MRG15 are seen in other mammalian species where MRG15 influences DNA repair, cellular differentiation, and pre-mRNA splicing in spermatogenesis (7, 8).

The structure of MRG15 consists of two conserved domains: an N-terminal chromodomain and a C-terminal MRG domain. The MRG15 chromodomain contains aromatic residues that form a barrel-like chromodomain pocket that interacts with modified histone tails, specifically H3K36me3 (9). When the chromodomain is mutated, MRG15 loses its transcriptional regulation activity suggesting that the chromodomain is necessary for its role in HAT activity(1). The MRG15 chromodomain barrel is conserved across multiple species including, mice, *Drosophila*, and *C. elegans*. In *Drosophila*, Mrg15 interacts with Ash1 and stimulates its H3K36 methyltransferase activity(10, 11). However, this interaction with histone methyltransferases is not known to be conserved in *C. elegans*(12).

Like MRG15, its *C. elegans* homolog MRG-1 has various functions, and has been found to co-purify with multiple chromatin modifying complexes. MRG-1 was recently identified as a member of the conserved HDAC complex, SINS3, that includes SIN-3 and HDA-1(12, 13). The SINS3 complex is thought to target histone deacetylation at different promoters, but the specific role of MRG-1 in this complex is unclear (13).

*C. elegans* MRG-1 is a maternally loaded protein that is essential for germline development. Maternally provided MRG-1 is both necessary and sufficient for the proper proliferation of primordial germ cells (PGCs) and the loss of MRG-1 results in improper regulation of germline-expressed genes (14–16). For example, MRG-1 is required for the suppression of numerous genes on the X-chromosome as well as the repression of germline-specific genes in somatic tissues (17). Conversely, MRG-1 also prevents ectopic expression of somatic genes in the germline, further confirming its role in transcriptional regulation in the germline(12). In addition to its transcriptional regulation functions, MRG-1 has also been shown to play roles in genomic integrity and meiosis. Mutants lacking MRG-1 exhibit a meiotic defect in which homologs are able to pair properly, but are unable to fully align for normal synapsis (18). The maternal effect sterility of *mrg-1* is similar to that seen in *mes-4* mutants. MES-4 is an H3K36 methyltransferase that, like *mrg-1,* has a maternal requirement that is both necessary and sufficient for germline development. The similarity between *mes-4* and *mrg-1* mutant phenotypes, and the known binding of H3K36me3 by MRG-1’s homolog MRG15, have led to the assumption that MRG-1’s essential germline role is through its interactions with MES-4-dependent H3K36me3 (19).

Here we further explore the role of MRG-1 in *C. elegans* meiosis and development. We confirm MRG-1’s essential function in meiosis and further characterize the synapsis defects in *mrg-1* mutants, including a role in proper crossover formation and regulation. We also show that a lack of zygotic MRG-1 protein causes a variety of partially penetrant somatic developmental defects. Given the presumed role of MRG-1’s chromodomain in H3K36me3 recognition, a modification with important germline functions, we investigated the specific roles of H3K36me3 and the chromodomain of MRG-1 in meiotic events. Depletion of H3K36me3 in the germline results in meiotic defects that are similar to absence of the MRG-1 protein. Surprisingly, we find that an intact chromodomain is not necessary for MRG-1’s function in meiosis or germline proliferation, but is instead required zygotically for embryonic development.

Oddly, mutation of the chromodomain causes novel phenotypes, including embryonic lethality and a dominant RNAi defect that is not observed in the null allele. This suggests that the roles of H3K36 methylation and MRG-1 in germline development and function are complex, and that MRG-1’s presumed recognition of H3K36 methylation may not be essential for post-embryonic germline development and meiotic regulation.

## Results

### Zygotic Requirements for MRG-1 in Meiosis

The human homolog of MRG-1, MRG15, plays important roles in transcriptional regulation(1–5). In *C. elegans,* MRG-1 is maternally loaded into the germline and the maternal contributions is required for proper germline development. MRG-1 is maternally necessary and sufficient for fertility: mutants have a grand- daughterless/maternal effect sterile (MES) phenotype in which homozygous *mrg-1/mrg- 1*mutants from heterozygous *mrg-1/+* mothers develop a germline, but the next generation is sterile. The fertile F1 progeny are termed M+Z-; i.e., they inherited maternal MRG-1 produced by the mother (M+), but lack a zygotically functional *mrg-1* gene (Z-). F2 progeny from M+Z*-* mutants are thus sterile because they do not inherit a maternal source required for germline development (M-Z-). Despite the *mrg-1* M+Z- offspring being fertile, they display a decreased brood size and increased embryonic lethality when compared to wild type or *mrg-1* heterozygotes (S1). This suggests that there is also a zygotic requirement for MRG-1 in the post-embryonic germline and soma.

We further confirmed results from previous studies that showed that zygotic MRG-1 is necessary for proper meiotic progression. In addition to increased sterility and embryonic lethality (S1), Carolyn et al. showed that *mrg-1* M+Z- offspring exhibit a homologous chromosome alignment defect: homologs can successfully pair at their pairing centers, but the chromosome ends opposite these centers fail to align (18). Homolog pairing and alignment occurs in the transition zone of *C. elegans* meiosis before progressing into pachytene and engaging in synapsis. The transition zone is characterized by nuclei with condensed DAPI-stained chromosomes grouped at one side of the nuclear periphery in a characteristic crescent shape. We compared the length of the transition zones in N2 wild type and *mrg-1(tm1227) M+Z-* mutants. In wild type hermaphrodites, the transition zone consists of a short region encompassing roughly 25-30% of the ovary region measured (described in Materials and Methods) (Fig 1A, C). In *mrg-1 M+Z-* worms, the transition zone length was significantly extended to as much as 50% of the calculated length and nuclei with the crescent-shape chromatin were also apparent in more proximal regions of the expected pachytene germline (Fig 1B, C). An extended transition zone is characteristic of delayed progression into pachytene and synaptic delay, as might be expected from previous reports of MRG-1’s role in pairing-center independent alignment. Therefore, despite having a maternal load of MRG-1 to initiate germline development, zygotic expression of MRG-1 is necessary for normal meiotic progression and synapsis (18).

**Figure 1.**
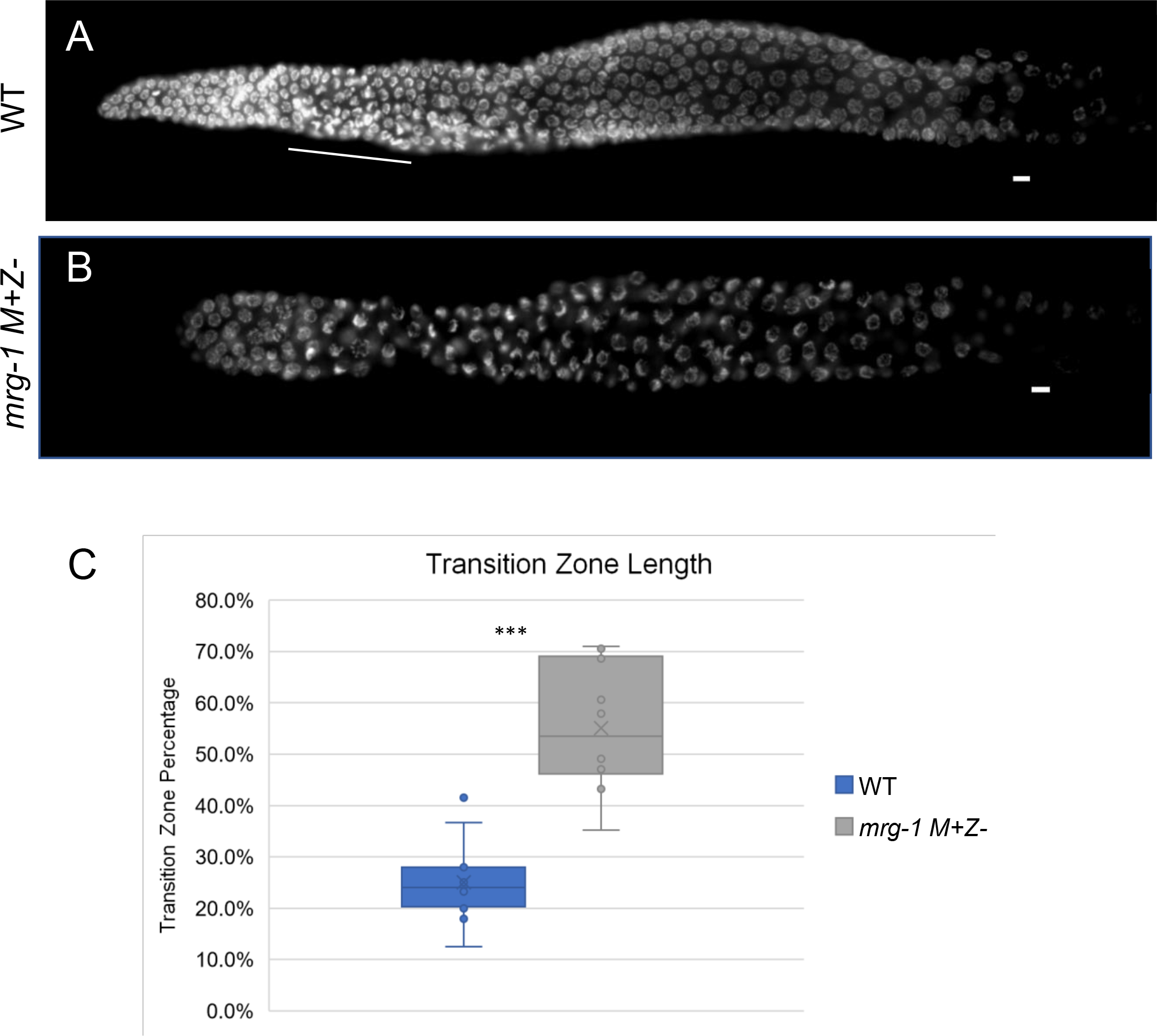
*mrg-1 M+Z-* mutants exhibit synaptic delay. Ovaries of WT and *mrg-1(tm1227) M+Z-* hermaphrodites stained with DAPI. The transition zone (white line) was measured in each germline and compared to the length from the distal tip to the end of pachytene as described in the Materials and Methods(C). In comparison to the WT, *mrg-1* mutants display an extended TZ. The total length of the TZ is 25% in WT germlines. In the *mrg-1* mutants, the TZ is over 50% of the determined length. Germlines were dissected, fixed, and stained with DAPI 24 hrs post L4 larval stage (*** indicate p ≤0.001).

### MRG-1 is required for the regulation of crossovers during meiosis

Given the defects in synapsis progression, we further characterized crossover formation in *mrg-1(tm1227)* mutants. In *C. elegans,* 6 pairs of homologous chromosomes must align, synapse, and initiate double-stranded breaks for recombination. One double strand break matures into a single crossover (CO) between each homolog that is marked by localization of the COSA-1 protein (20). In worms expressing GFP::COSA-1, 6 GFP foci are visible in late pachytene nuclei (region 6: n=102) where 6 COs have successfully formed between homologs (Fig 2A, C). Upon *mrg-1* knockdown by RNAi, 6 COSA-1 foci were observed in the majority of nuclei (region 6: n=63), yet nuclei with less than 6(region 6: n=28) and some with more than 6 COSA-1 foci (region 6: n=8) were also often observed, indicating a defect in CO regulation (Fig 2B, D). Additionally, CO formation appears to occur earlier in pachytene (Region 4) after *mrg-1* knockdown. As cross-over interference and maturation is regulated by proper synaptonemal complex formation between homologs, this is likely another downstream effect of defective homolog alignment and abnormal synapsis with MRG-1 knock-down(21).

**Figure 2.**
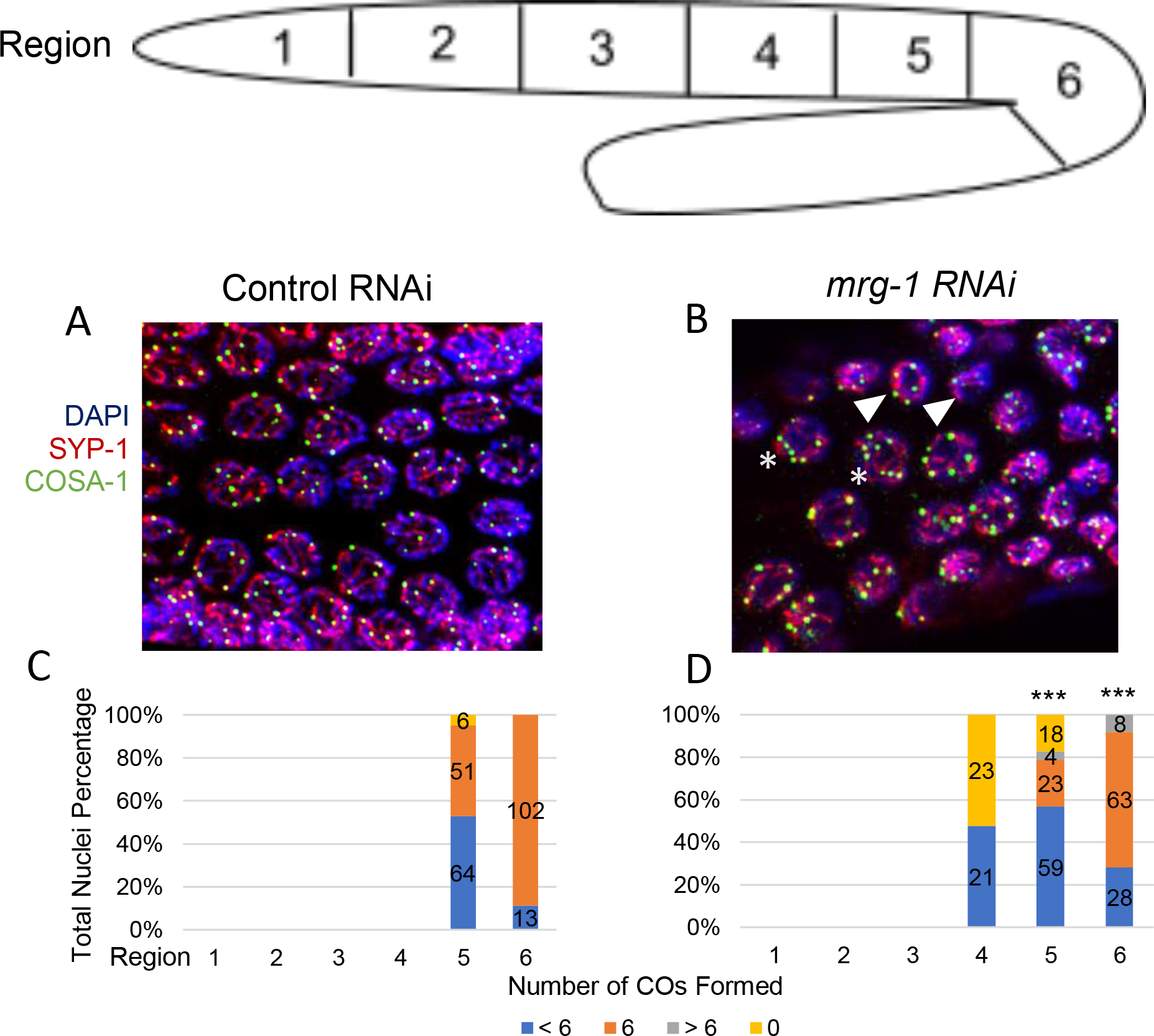
*mrg-1 M+Z-* mutants have defective crossover regulation. Ovaries from control (A) or *mrg-1(RNAi)* (B) treated animals were dissected, fixed, and stained with anti-COSA-1, anti-SYP-1, and counterstained with DAPI. The total length of the gonad was divided into 6 regions and COSA-1 signals per nuclei were counted in each region (C, D). At the end of pachytene (Region 6), WT nuclei (C) have 6 mature crossovers (CO) marked by COSA-1 for each set of homologs. In *mrg-1* mutants (D), CO formation often occurs earlier in pachytene and have an abnormal number of COs (arrowhead <6, asterisk >6) (*** indicate p ≤0.001).

As expected for defects in synapsis and CO formation, we also observed an increased number of achiasmatic chromosomes in diakinetic oocytes. In wildtype oocytes, 6 attached pairs of homologs (bivalents) are normally observed in the oocytes (n=121)(Fig 3A). In contrast, *mrg-1(tm1227) M+Z-* mutants exhibit a variable number of univalent chromosomes(n=25), with as many as 12 DAPI bodies observed in some oocytes (Fig 3B). It is important to note, however, that at least 50% of the *mrg-1* oocytes successfully synapsed and recombined all homolog pairs(n=36), indicating that zygotic MRG-1 is only partially required for proper homolog alignment, synapsis, and recombination (Fig 3C).

**Figure 3.**
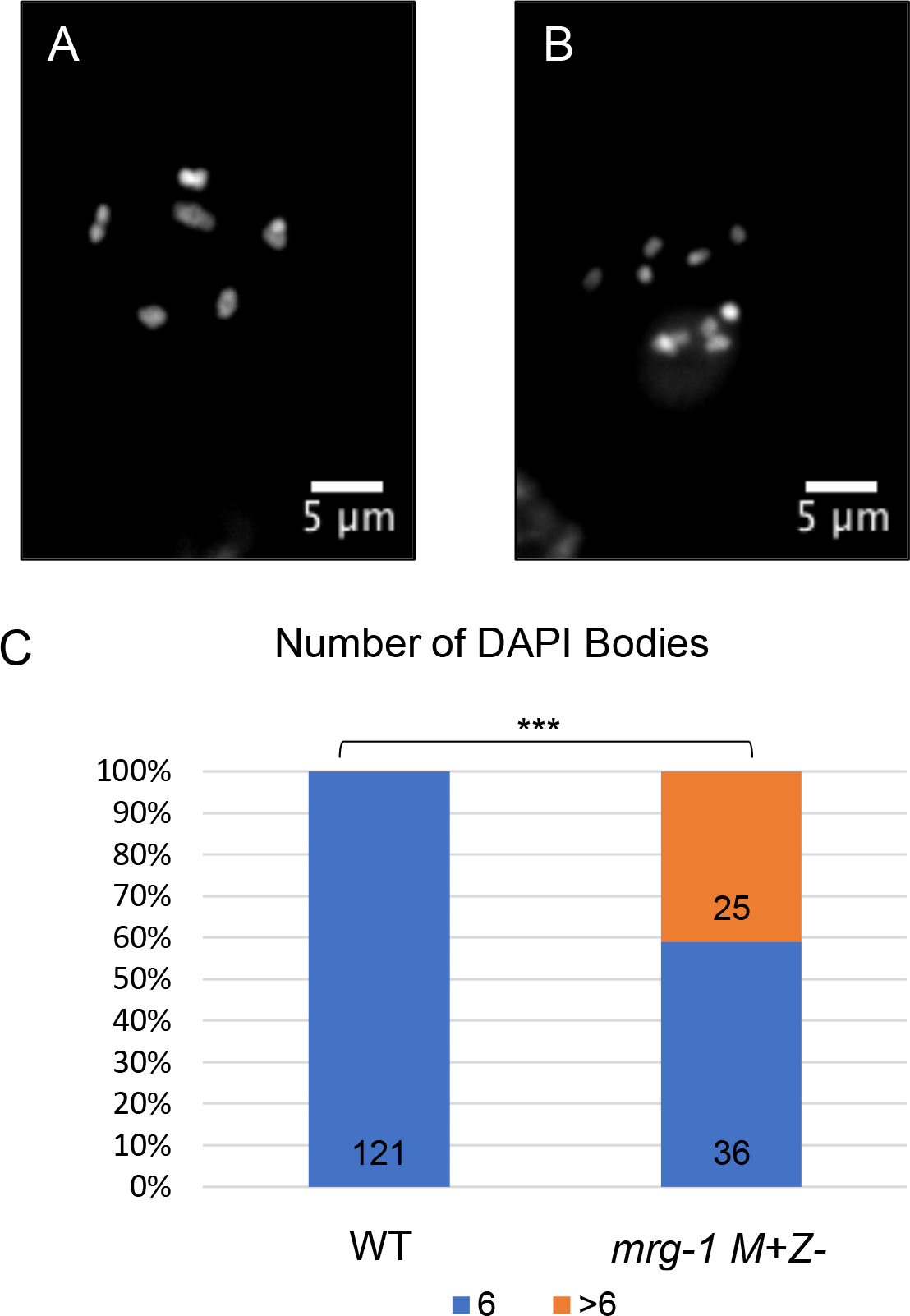
*mrg-1 M+Z-* mutants have an increased number of achiasmatic chromosomes. (A) Oocytes in wildtype germlines have 6 bivalents that are generated after homologs have successfully recombined and formed productive crossovers. (B) Mutants lacking zygotic MRG-1 display a range in number of DAPI bodies suggesting that recombination is defective (C). Germlines were dissected, fixed, and stained with DAPI 24 hours after L4 larval stage (*** indicate p ≤0.001).

### H3K36me3 is Required for Normal Meiotic progression

As a homolog of MRG15, it is predicted that MRG-1 recognizes and binds to H3K36 methylation. Indeed, ChIP analyses of MRG-1 has shown its enrichment in the genome overlaps with enrichment of H3K36me3 and H3K4me3 (12). In order to determine if H3K36me3, and thus its recognition by MRG-1, were associated with the *mrg-1(tm1227)* phenotypes, we reduced H3K36 methylation in germ cells and assessed any meiotic consequences. MET-1 and MES-4 are the two H3K36 methyltransferases in *C. elegans*. MET-1 dependent H3K36 methylation is associated with active transcription, whereas MES-4 activity can occur independently of transcription and is thought to play a role in maintaining an epigenetic memory of germline transcription across generations (22–24). Single RNAi knock downs, and/or single mutations of either *mes-4* or *met-1* do not substantially decrease H3K36me3 levels in germ cells (Fig 4 A, B, E). Previous reports demonstrated that the disruption of both *met-1* and *mes-4* depletes all H3K36me3 in embryos (22). We therefore performed *mes-4* RNAi in L1 *met-1* mutant larvae, which had dramatically reduced H3K36me3, and analyzed their germlines (Fig 4F).

**Figure 4.**
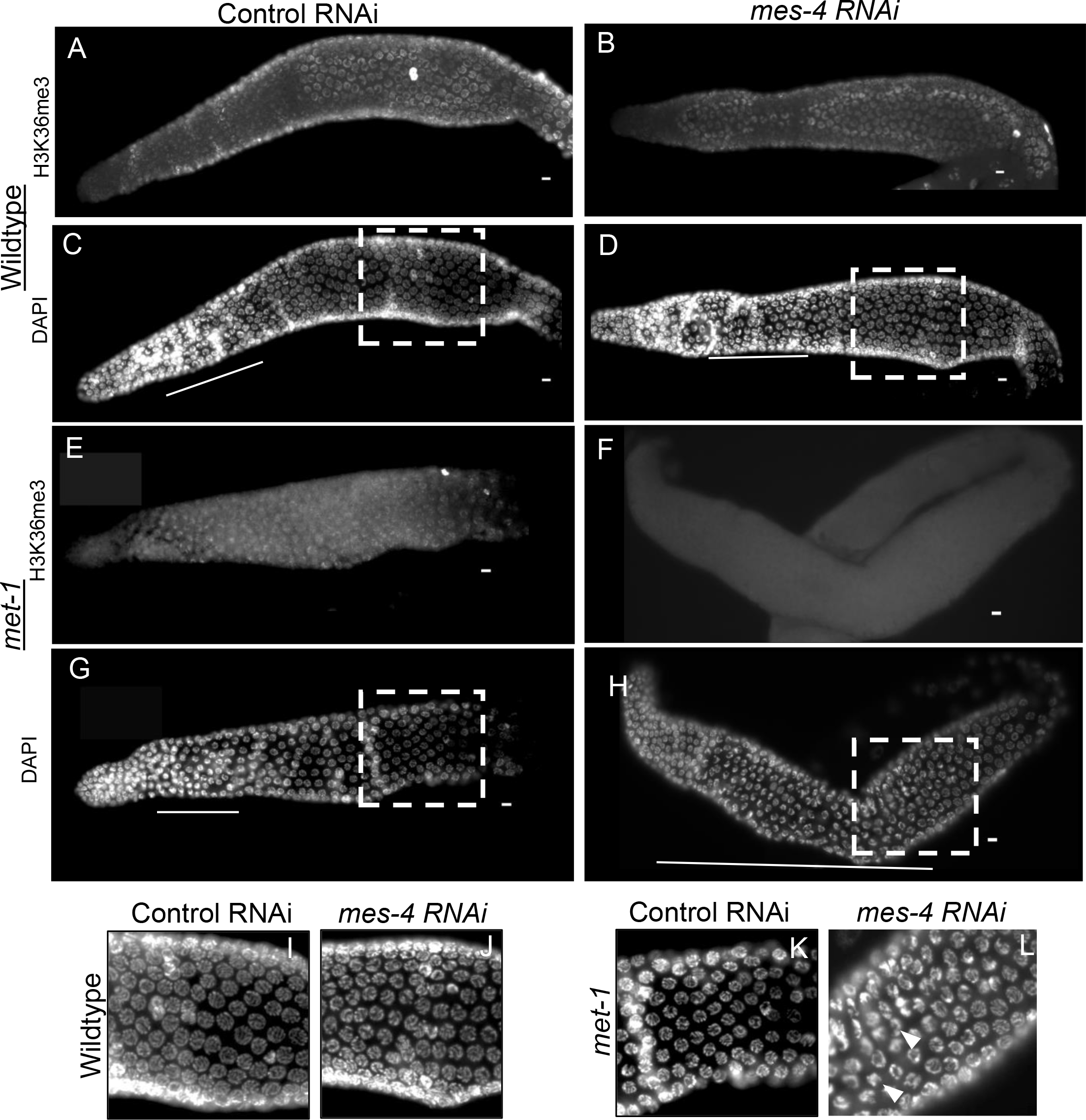
Depletion of H3K36me3 causes synaptic delay. Ovaries from wildtype (A, C), *mes-4(RNAi)* (B, D), *met-1(n4337)* (E, F), and *met-1(n4337); mes-4(RNAi)* adult hermaphrodites were dissected, fixed, and probed with anti-H3K36me3 followed by DAPI staining. H3K36me3 is present in A, B, and E, but is reduced in F. The transition zone (indicated by white line) is restricted to the distal gonad in A, D, and G, but is highly extended in H. Panels I-L are enlargements of regions outlined in panels C, D, G, and H. Arrowheads indicate nuclei with condensed chromatin located at the periphery, similar to those observed in the transition zone.

We first asked if reduced H3K36me3 results in an extended transition zone, as observed in *mrg-1* M+Z- ovaries. In wildtype germlines, after being treated with both the empty control vector and *mes-4* RNAi, normal lengths of synapsis were visible and no transition zone nuclei were present in the proximal pachytene region (Fig 4C, D). A similar result was seen in *met-1(n4337)* and in control RNAi germlines (Fig 4G) suggesting that loss of either MET-1 or MES-4 activities alone has little effect on meiotic progression. In contrast, *met-1(n4337); mes-4(RNAi*) worms exhibited crescent shaped nuclei visible throughout the germline extending into the proximal pachytene region (Fig 4H), indicating a delay in synapsis. This extended transition zone phenotype suggests that H3K36me3, like MRG-1, plays an important role in meiotic progression.

### M-Z- *mrg-1* animals Exhibit Zygotic Defects in Soma

Maternal deposition of MRG-1 is essential for fertility and M-Z- offspring from fertile M+Z- mothers are sterile but grow into adulthood. However, we found that the M- Z- generation showed a percentage of the worms that reach adulthood displayed somatic developmental defects. *mrg-1(tm1227)* M-Z- animals frequently displayed somatic defects such as multiple vulvas (Muv), tail defects, and dumpy (Dpy) morphology (S Table 1). These somatic phenotypes in the M-Z- generation indicate that zygotic expression of MRG-1 is also important for normal somatic development. (S2)

### Mutations in MRG-1’s Chromodomain Cause Embryonic Lethality

Our results from depleting H3K36me3 from germ cells implied that the role of MRG- 1 in meiosis is linked to its presumed role as a conserved “reader” of H3K36 methylation. In addition, the somatic defects observed in *mrg-1(tm1227)* M-Z- animals suggested a role for zygotic MRG-1 in somatic development, which may also be related to its presumed role in H3K36me3 recognition. To investigate this, we mutated amino acids in two conserved positions that in MRG15 form the aromatic cage of the chromodomain, Y17 and F43, to alanines and assessed their effects on germline and somatic phenotypes. We first generated both GFP and mCherry tagged versions of the endogenous wild-type (WT) *mrg-1* gene using CRISPR. Both strains carrying the WT endogenous tagged versions had normally fertility and exhibited no overt phenotypes (below). We then further replaced Y17 and F43 with alanines in the *mrg-1::mCherry* strain in another round of CRISPR, yielding a new allele, *ck43,* which we will hereafter refer to as *mrg-1^CD^* (Materials and Methods).

As mentioned, maternal provision of MRG-1 protein is necessary for germline development, and the M+Z- generation of *mrg-1(tm1227)* is fertile. The M-Z- offspring of homozygous *mrg-1(tm1227)* animals are viable but fail to develop a germline. After verifying and outcrossing the *mrg-1* chromodomain mutant (*mrg-1^CD^*), it was genetically balanced with the qC1 chromosome for reasons described below. Although the *mrg- 1^CD^/qC1* exhibited decreased brood sized when compared to wildtype/*qC1*, they were fertile. Surprisingly, 100% of the *mrg-1^CD^/mrg-1^CD^* offspring from heterozygous mothers died as embryos (S3, Fig 5). Therefore, whereas the *mrg-1(tm1227)* null allele causes maternal effect sterility but no lethality, the *mrg-1CD* mutation causes zygotic lethality despite inheriting maternal WT MRG-1 from the qC1 balancer chromosome. This lethality occurs as a mid-embryogenesis arrest: the arrest point appeared somewhat variable with some embryos developing to where movement was observed, but others did not. The *mrg-1CD* mutant’s embryonic lethal phenotype is therefore worse than the homozygous null phenotype, which survives embryogenesis in the absence of any MRG-1 protein. The presence of one dose of maternal MRG-1 protein in the *mrg- 1^CD^/mrg-1^CD^* offspring from heterozygous mothers doesn’t overcome the effect of the CD mutation, suggesting this allele is acting dominantly, or that the maternal load produced from a single copy of the WT gene is insufficient to overcome the deleterious effects of the CD mutation (S7). The normal development of *mrg-1^CD^/+* animals suggests that the phenotype is more complicated than a dominant or antimorphic situation, although a non-lethal yet dominant phenotype *is* observed in these animals (described below).

**Figure 5.**
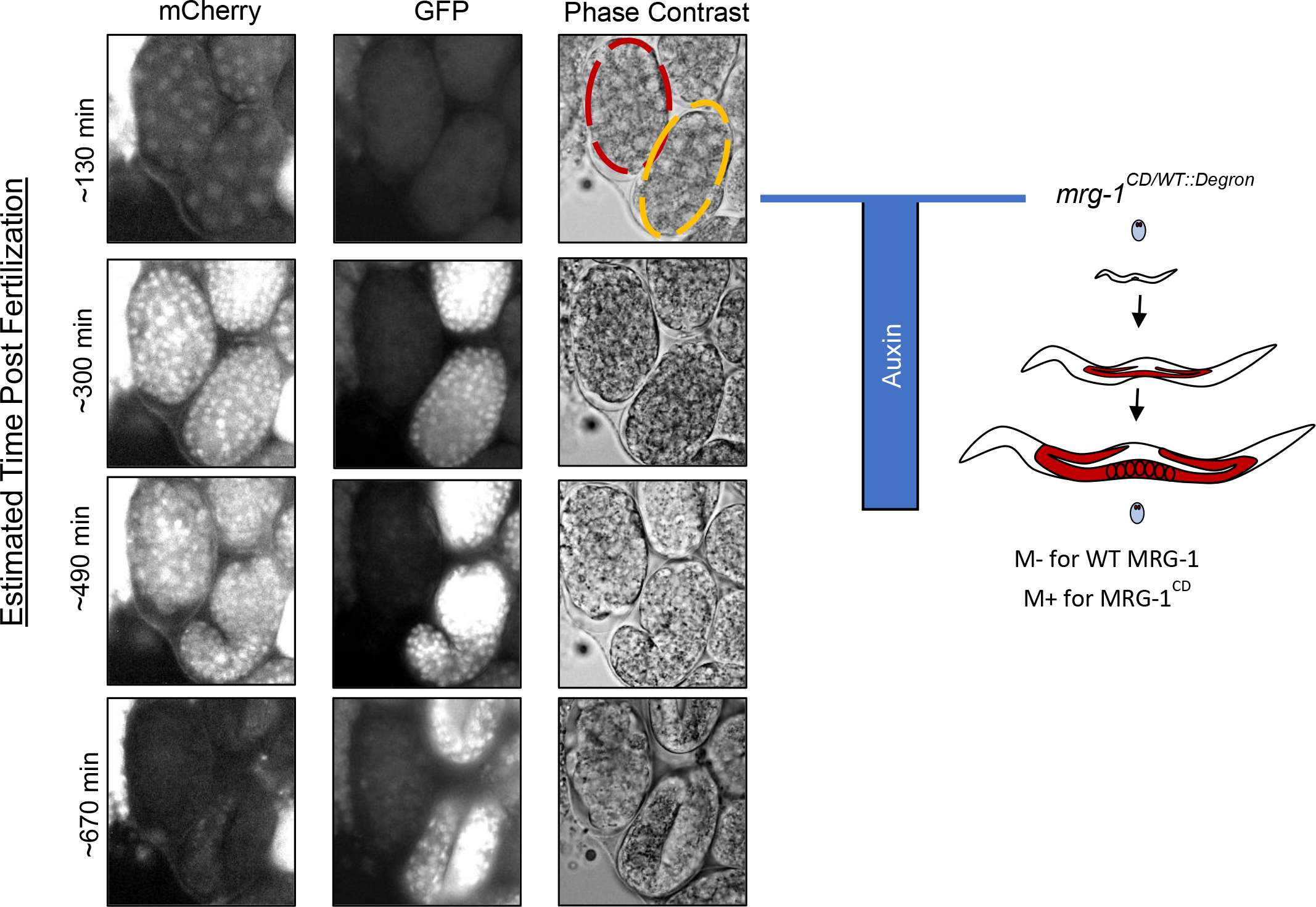
*mrg-1* Chromodomain Mutants arrest during embryogenesis. Embryos were dissected from *mrg-1^CD/WT::Degron^* hermaphrodites exposed to auxin beginning at hatching and analyzed by fluorescence and phase microscopy over eleven hours. Embryos expressing zygotic GFP, either homozygous *mrg-1^WT::Degron^* or *mrg- 1^CD/WT::Degron^* (yellow outline) developed normally. In contrast embryos expressing only *mrg-1^CD^* (red outline) arrested in early- to mid-embryogenesis.

### Maternally provided CD mutant MRG-1 alone is not sufficient for germline development

Because the homozygous *mrg-1^CD^* mutants die as embryos, we utilized the Auxin Inducible Degradation (AID) System to further examine the role of the chromodomain in the post-embryonic germline. In order to bypass the embryonic lethal phenotype, we used CRISPR to add the AID degron to the endogenous (WT) *mrg-1* locus tagged with GFP, producing an *mrg-1::degron::gfp* strain. We crossed the *mrg-1::degron::gfp* strain into a strain carrying TIR1 driven by the *sun-1* promoter for germline specific degradation (*mrg-1*^WT::Degron^). We confirmed that auxin induced degradation of the MRG-1::degron::GFP protein in adults produced sterile offspring, similar to M-Z- *mrg- 1(tm2337)*. We then crossed the *mrg-1^CD^ /qC1* balanced line with the *mrg-1*^WT::Degron^ to generate *mrg-1*^CD/WT::Degron^ heterozygous animals. In *mrg-1*^CD/WT::Degron^ heterozygous animals, MRG-1^CD^ mCherry and MRG-1^WT::Degron^ GFP were visible and colocalized in the germline (S5). The addition of the AID degron allowed controlled degradation of the wildtype MRG-1^WT::Degron^, leaving only the mutant protein present and allowing us to examine the functional consequences of the CD mutations. Because TIR1 is driven by the germline-specific *sun-1* promoter, exposing the MRG-1^CD/WT::Degron^ animals to auxin yielded germline-specific degradation of MRG-1^WT::Degron^, leaving only the MRG-1^CD^ protein (detectable by its mCherry tag) present in the germline. As auxin is reported to poorly penetrate the embryo’s eggshell (25) and thus unlikely to activate WT degradation via any maternal TIR1, the embryos bypass the CD mutant’s embryonic lethality through zygotic expression of the WT protein in the embryos’ soma (26). When *the mrg-1^CD/WT::Degron^* embryos hatch onto auxin-treated plates, the WT protein is continuously degraded in the post-embryonic germline, allowing separation of somatic and germline requirements for an intact chromodomain to be assessed (Fig 5).

Degradation of the MRG-1^WT::Degron^ by auxin treatment was confirmed visually by the absence of GFP in the germline. Additionally, embryos with a maternal load of MRG-1^WT::degron^ that were exposed to auxin upon hatching showed immediate degradation of maternal WT MRG-1^WT::Degron^ and grew up sterile mimicking the MES phenotype (Fig 7). Previous studies have shown that the *mrg-1* promoter is not active in the embryonic germline (27), thus at the earliest post-hatching stages maternal protein may be initially still necessary for fertility but also immediately susceptible to auxin degradation. It is important to note that no noticeable degradation of MRG-1^WT::Degron^ was observed when animals were grown without auxin suggesting there was no nonspecific activation of TIR1.

**Figure 6.**
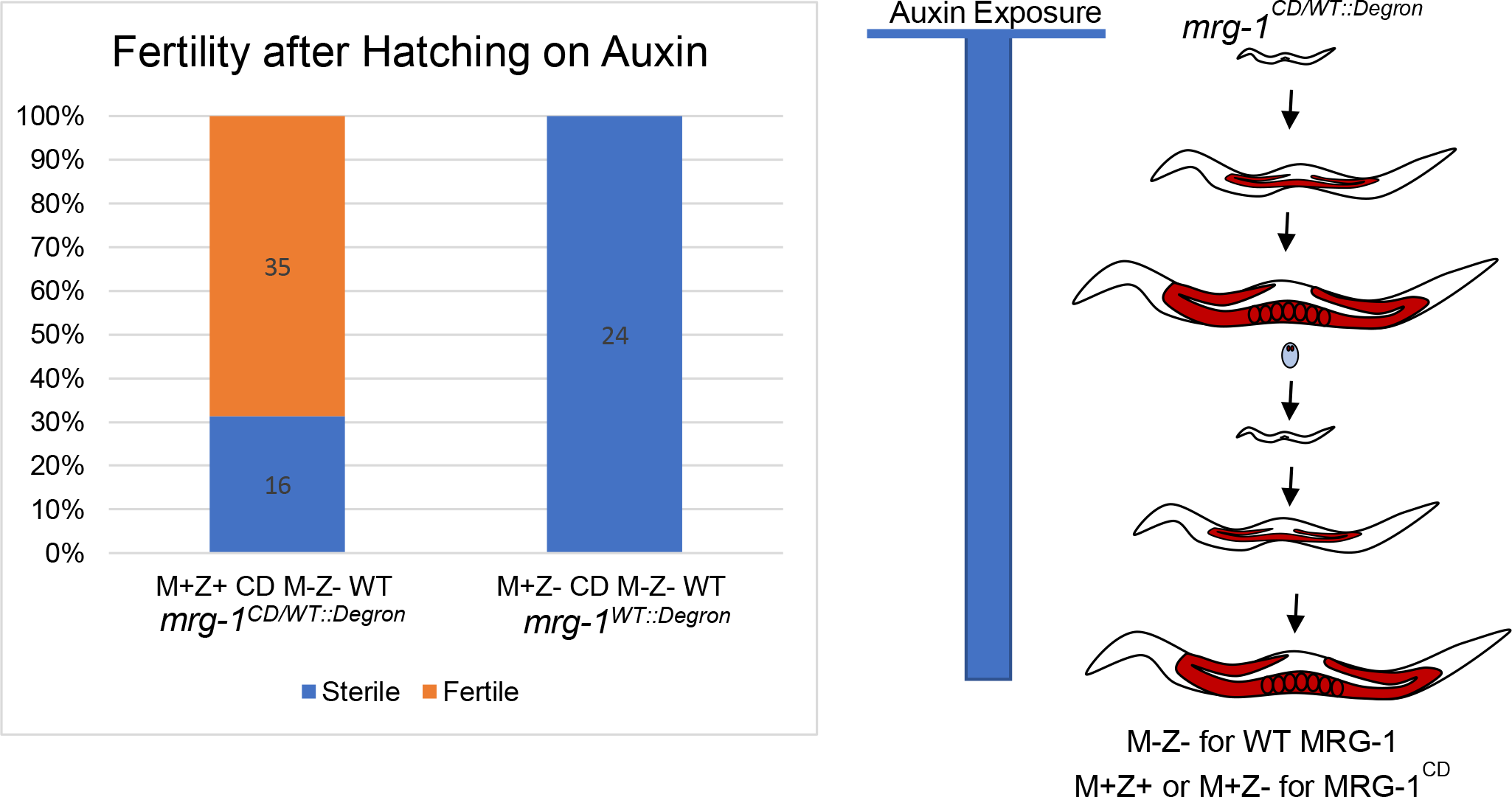
Maternal MRG-1 Chromodomain mutant alone is not sufficient for germline development. Embryos dissected from MRG-1^CD/WT::Degron^ hermaphrodites exposed to auxin during all of larval and adult development were hatched on auxin plates, yielding offspring that were M-Z- for WT MRG-1, all of which inherited maternal CD mutant, and 2/3 of which also expressed the CD mutant zygotically in their post- embryonic germline. All offspring M-Z- for WT MRG-1 grew up sterile. In contrast, 75% inheriting maternal *mrg-1*CD in combination with zygotic *mrg-1CD* mutant grew up fertile.

**Figure 7.**
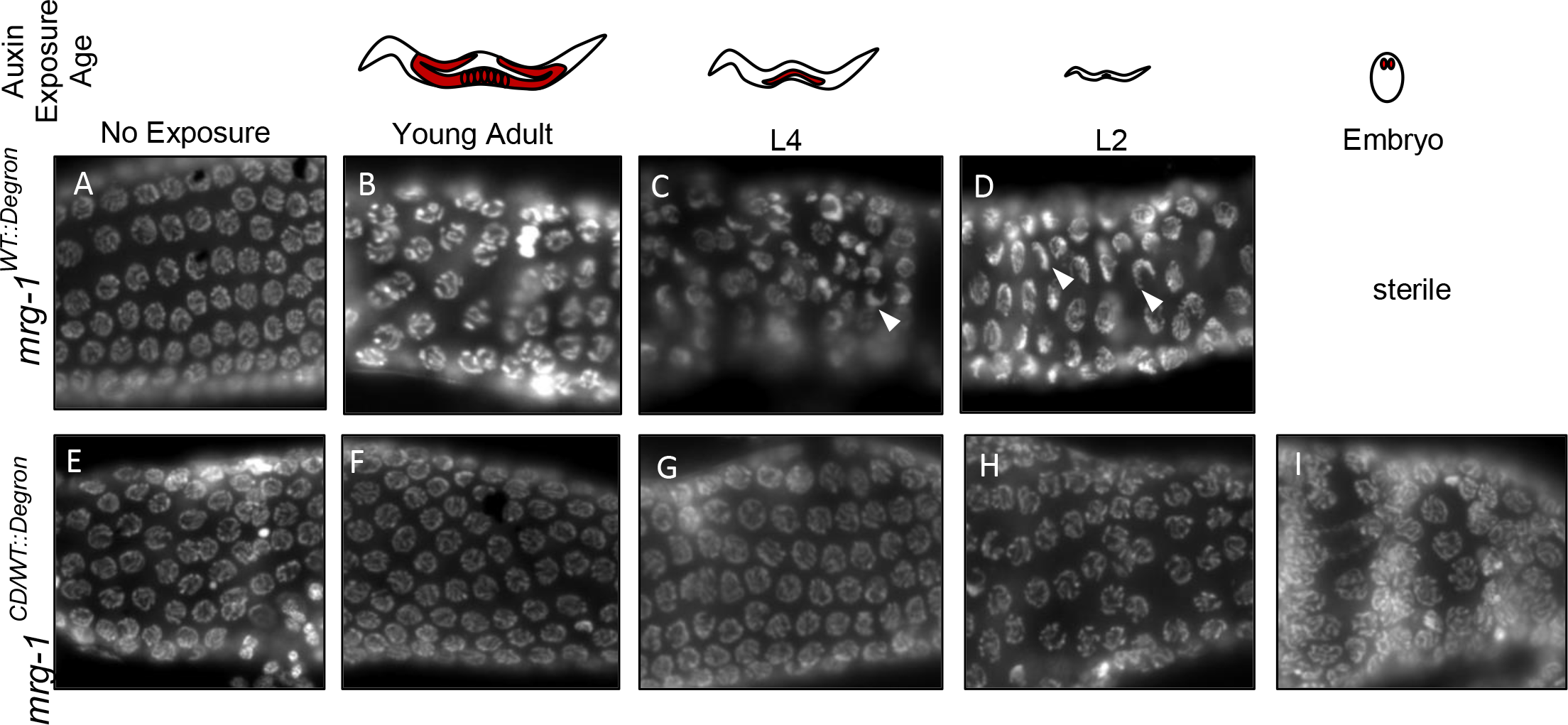
MRG-1 Chromodomain mutant supports normal meiosis. *mrg-1^WT::Degron^* (A-D) and *mrg-1^CD/WT::Degron^* (E-I) worms were placed on auxin (1mM) starting from various larval stages (indicated by blue bar). Gonads dissected from adult animals from each treatment were then fixed and stained with DAPI. The proximal region where synapsis normally occurs is shown. Nuclei exhibiting pre-synaptic delay are indicated by a crescent-like chromosome organization. Auxin-induced degradation of *mrg-1^WT::Degron^* starting from L2 larvae caused highly penetrant synaptic delay (arrowheads; C-D), whereas auxin exposure starting in adult animals caused chromosome morphology defects (B) but had minimal effect on synapsis. Earlier auxin degradation of WT MRG-1 begun at hatching caused sterility with no germline. In contrast, germlines with only the CD mutant present beginning at any stage showed no overt meiosis defects (E-I).

To determine if a functional MRG-1 chromodomain was required for germline development, embryos from *mrg-1*^CD/WT::Degron^ hermaphrodites exposed to auxin were hatched on auxin plates. The heterozygous offspring therefore lacked both maternal and zygotic WT MRG-1 in their germ cells, with only the MRG-1^CD^ mutant protein present in their germline at all stages (M-Z- for WT MRG-1, M+Z+ for MRG-1^CD^) (Fig 6). Their *mrg-1*^WT::Degron^ homozygous siblings had inherited only maternal MRG-1^CD^ protein, and any WT protein was degraded upon hatching. Individual animals were observed under a microscope to determine fertility status and genotyped. 100% (n=24) of the homozygous MRG-1^WT::Degron^ offspring from auxin-exposed mothers were sterile. This suggests that the maternal load of MRG-1 CD provided by their heterozygous mothers alone is on its own insufficient for germline development. In contrast, 69%(n=35) of the MRG-1^CD/WT::Degron^ were fertile and 31% were sterile (n=16) suggesting that maternal MRG-1 CD when combined with zygotic expression of *mrg-1*^CD^ is able to partially rescue germline development. As discussed above, the *mrg-1*^CD^ */ mrg-1*^CD^ homozygous offspring failed to hatch.

### The MRG-1 Chromodomain is not required for normal meiotic progression

Because many of the *mrg-1*^CD/WT::Degron^ embryos hatched on auxin were fertile and successfully bypassed the embryonic lethal phenotype, we were able to analyze germlines that only contained the mutated version of the MRG-1 chromodomain.

Because synapsis defects were apparent after reducing levels of H3K36me3 in the germline, we wanted to determine if similar phenotypes were seen after mutating the MRG-1 chromodomain that correlated with its presumed binding specificity. We placed *mrg-1*^WT::Degron^ and *mrg-1*^CD/WT::Degron^ on auxin plates starting from various larval stages and analyzed them as adults. When placed on auxin starting from L2 larval stage, *mrg- 1*^WT::Degron^ worms exhibited meiotic defects similar to the M+Z- *mrg-1(tm1227)* mutants, including delayed synapsis (Fig 7 C, D). The synapsis defects became less severe in *mrg-1*^WT::Degron^ worms if auxin degradation started after the young adult stage, despite having complete degradation of wildtype MRG-1 for 36 hours (Fig 7B*). mrg-*1^WT::Degron^ M*+Z- e*mbryos that hatched on auxin plates grew up to be sterile. This we assume to be due to the maternal load being immediately degraded shortly after hatching and before post-embryonic germline development, confirming a requirement for maternal, and possibly also zygotic, MRG-1 at the initiation of germline development.

*mrg-1^CD^*^/WT::Degron^ heterozygotes (Fig 7 E-I), were similarly exposed to auxin for various amounts of time from different larval stages. No synapsis defects were observed after any length of auxin exposure (Fig 7 F-I). Unlike the *mrg-*1^WT::Degron^ germlines that showed signs of delayed synapsis after auxin exposure from L2 larval stage, germlines with only the CD mutant appeared to have normal synapsis.

Additionally, *mrg-1*^CD/WT::Degron^ embryos that were hatched on auxin developed to be fertile, unlike their homozygous *mrg-*1^WT::Degron^ siblings that were sterile (Fig 7I). The apparently normal synapsis in *mrg-1*^CD/WT::Degron^ germlines after auxin exposure suggests that the chromodomain of MRG-1 is not important for MRG-1’s role in meiosis, in contrast to the MRG-1 protein itself. Therefore, MRG-1’s meiotic function may be dependent on a different region of the protein, its presence in a complex that is not dependent on the chromodomain for its function in the germline, or a redundancy for its role in meiosis. Complexes containing MRG-1may depend on MRG-1 protein for complex stability since the absence of MRG-1 protein causes significant germline defects.

### Zygotic MRG-1 combined with Maternal MRG-1CD partially rescues germline development

Previous studies have shown that the *mrg-1* promoter is not active in the embryonic germline, and thus all embryonic germline functions for MRG-1 must strictly rely on its maternal supply (27). To further characterize the maternal requirements for MRG-1 in the germline, we looked at embryos from *mrg-1^CD^*^/WT::Degron^ auxin-treated mothers that were hatched on OP50 bacteria plates lacking auxin. The offspring from these animals therefore had only the CD mutant as their maternal load, but zygotically expressed either both the CD and WT versions, or only WT *mrg-1* in their post-hatching germlines. Homozygous *mrg-1^CD^* embryos did not hatch, as described above (Fig 5). F1 offspring of the auxin treated mothers that were hatched without auxin were examined 24 hours after L4 for fertility and genotyped. Of the worms that were maternally supplied with CD and only zygotic for WT (M+Z- CD; M-Z+ WT) 43% were fertile (n=9) and 57% were sterile (n=12). This result suggests that zygotic expression of WT MRG-1 can only partially rescue germline development when coupled with maternal CD (Fig 8). Similarly, 53% (n=26) of offspring that had maternal CD and expressed both WT and CD zygotically (M+Z+ CD; M-Z+ WT) were fertile, while 47% (n=23) were sterile, again suggesting that zygotic expression of CD mutant and/or WT MRG-1 only partially rescues germline development when the maternal supply is the CD mutant. We note that some sterile worms that were M+Z+ CD; M-Z+ WT had partially developed germlines while others completely lacked any germ cell proliferation, similar to the maternal effect sterile phenotype (S4).

**Figure 8.**
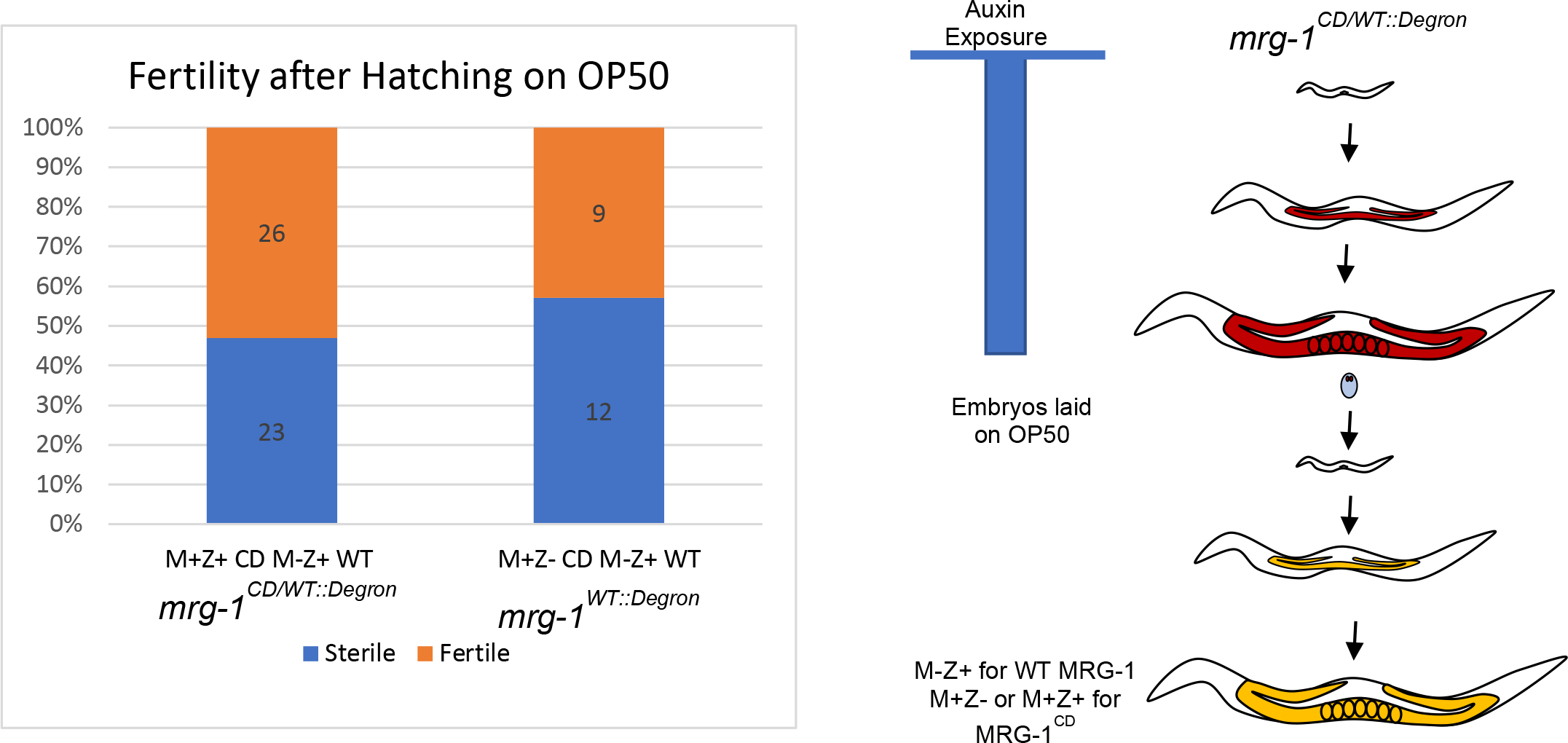
Zygotic expression of WT MRG-1 in combination with maternal MRG- 1CD can partially rescue fertility. Embryos from *mrg-1^CD/WT::Degron^* hermaphrodites exposed throughout post-embryonic development were hatched on OP50 plates and scored for fertility as adults. These worms inherited only maternal CD mutant MRG-1 but could express zygotic WT MRG-1 when hatched onto OP-50. Of the worms that had zygotic expression of WT, 40% were able to develop a germline. The presence or absence of zygotic CD mutant MRG-1 did not appear to impact fertility.

### The MRG-1 Chromodomain is necessary for exogenous RNA interference

We originally tried to use RNAi to target the GFP-tagged WT in *mrg-1::gfp/mrg- 1CD::mCherry* animals to bypass the embryonic lethality of the CD mutation and examine CD mutant germlines. However these experiments failed because we observed a striking RNAi resistant phenotype. As expected, GFP RNAi successfully eliminated MRG-1 in *mrg-1::gfp* homozygous worms, resulting in the absence of GFP in the germline and the expected MES phenotype in the next generation (Fig 9F).

**Figure 9.**
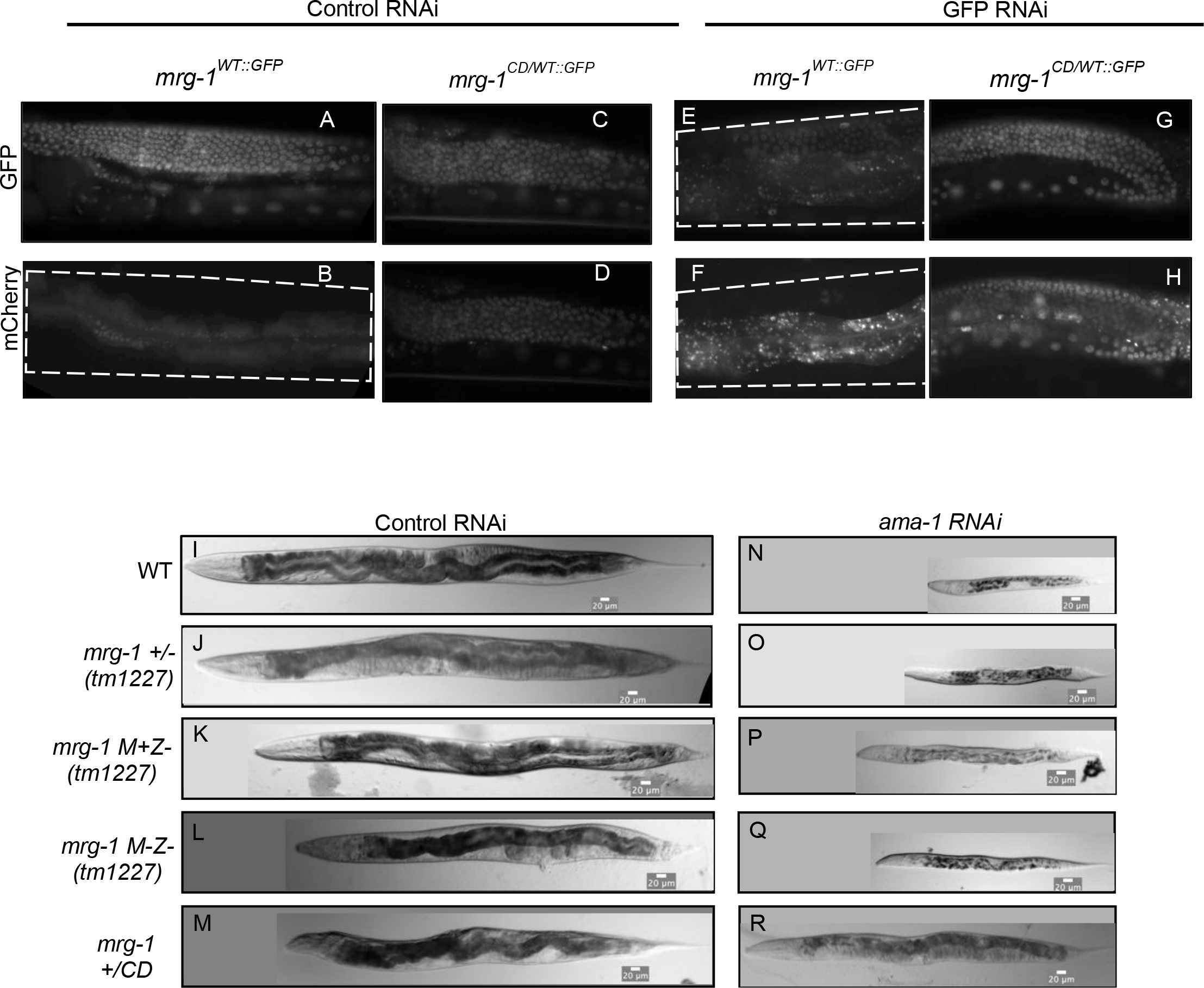
*mrg-1 CD* mutation causes a dominant RNAi resistance phenotype. WT MRG-1^WT::GFP^or heterozygous MRG-1^CD/WT::GFP^were placed on control(L4440) (A-D) or GFP RNAi (E-H) plates as L1 larvae. GFP RNAi depleted the GFP signal in MRG-1^WT::GFP^ animals, but was ineffective in animals carrying the CD mutant allele. (B) Wildtype, *mrg-1/+ (tm12227), mrg-1 (tm1227) M+Z-, mrg-1 (tm1227) M-Z-,* and *mrg-1 CD/+* worms were placed on control (I-M) or *ama-1 RNAi* (N-R) plates as L1 larvae and grown for >48 hours. All strains except *mrg-1 CD/+* (R) displayed L1 larval arrest.

However, we found we could not deplete the GFP signal by GFP RNAi in *mrg- 1::gfp/mrg-1CD::mcherry* heterozygous animals, suggesting that the presence of the *mrg-1 CD* mutation created a dominant resistance to the GFP RNAi (Fig 9H).

To further confirm this was an RNAi defect, we performed RNAi knockdown of *ama-1* in heterozygotes for the chromodomain mutant as well the *mrg-1(tm1227)* deletion allele. RNAi targeting *ama-1* (the gene encoding the large subunit of RNA polymerase II) in wildtype L1 larvae causes an arrested growth phenotype, and the larvae do not develop past L1-L2 stage (Fig 9N). This arrest phenotype is also seen in *mrg-1(tm1227) M+Z-* and *mrg-1 (tm1227) M-Z-* null mutants exposed to *ama-1(RNAi)* (Fig 9O-Q). However, no arrest in larval development was observed in *mrg-1CD/mrg- 1WT* L1’s exposed to *ama-1(RNAi)* (Fig 9R). Chromodomain mutants display a slight reduction in length when compared to the RNAi control but otherwise developed into fertile adults.

RNAi experiments targeting animals to *him-3(RNAi)* yielded similar results: the CD/WT animals exhibited resistance to RNAi, whereas the other genotypes did not (S6). We then examined whether CD/CD animals were RNAi defective using the AID system described above. Animals in which the WT MRG-1 had been degraded in their germlines with only the CD mutant present were also defective in RNAi (data not shown).

These results indicate that the CD mutation confers a dominant, gain-of-function phenotype on the MRG-1 protein that leads to an RNAi defective phenotype in both somatic lineages and the post-embryonic germline. MRG-1’s chromodomain is thus essential for somatic development in the embryo and its function is important for exogenous RNAi, but it is not essential for post-embryonic germline development. This indicates there are separable roles for MRG-1 and its chromodomain in somatic and germline development.

## Discussion

### Both maternal and zygotic MRG-1 are required for normal germline development and function

As a homolog of MRG15, it is expected and indeed has been shown that MRG-1 plays various roles in *C. elegans* development. Here, our goal was to further characterize MRG-1’s role in germline development, its presumed connection with H3K36me3 in germline function, and the role of its conserved chromodomain in its germline function. Consistent with previous reports, our data demonstrates that in addition to its maternal requirement, zygotically produced MRG-1 is also necessary for proper germline function and normal meiosis in *C. elegans.* Worms that have only a maternal load and no zygotic expression of *mrg-1* (M+Z-) display significant meiotic defects despite inheriting the maternal protein, including synaptic delay and defective crossover formation. Zygotic MRG-1 is thus required for normal meiotic progression and synapsis, consistent with previous studies by Carolyn et al. showing that zygotic MRG-1 is necessary for pre-synaptic alignment (18).

As MRG-1 is a proposed reader of H3K36me3, we investigated whether depleting H3K36me3 in the germline would yield consequences similar to the depletion of zygotic MRG-1 in the *mrg-1(tm1227)* M+Z- animals. Germlines that lacked normal levels of both MET-1 and MES-4, the two histone methyltransferases responsible for H3K36me3, exhibited efficient depletion of H3K36me3 and delayed meiotic progression and synapsis similar to that observed in *mrg-1* M+Z- mutants, suggesting that normal levels of H3K36me3 in meiotic chromatin are required for meiotic processes. Indeed, MET-1 and MES-4, and thus presumably the H3K36 methylation they provide, have been shown to be required for checkpoint activation(28). This result is also consistent with a similar report that looked at the contributions of H3K79 methylation in meiotic progression(29). Although it is unclear how specific histone modifications are mechanistically linked to meiotic processes, it is not surprising that chromatin organization plays an important role, given the unique architecture of meiotic chromatin.

Our data also further emphasizes the importance of the maternal load of wildtype MRG-1 in the early, post-embryonic germline. The promoter for MRG-1 is not active in the embryonic germline, and thus all MRG-1 present in the primordial germ cells (Z2 and Z3) before hatching is maternally supplied (27). In MRG-1^WT::Degron^ homozygote embryos that hatch on auxin, the immediate degradation of the maternal load in their germline yields 100% sterility, similar to the maternal effect sterility seen in the *mrg- 1(tm1227)* deletion mutant. Importantly, in MRG-1^WT::Degron^ embryos from *mrg- 1*^CD/WT::Degron^ + auxin mothers (M-Z+ WT; M+Z- CD) that hatch on OP-50, fertility is only partially rescued by maternal CD mutant MRG-1 when in combination with early zygotic WT MRG-1. A percentage of the sterile MRG-1^WT::Degron^ (M-Z+ WT; M+Z- CD) show germ cell proliferation, but do not fully rescue fertility. In contrast, homozygous *mrg*- 1^WT::Degron^ embryos from *mrg*-1^CD/WT::Degron^ + auxin mothers that hatch on auxin, therefore only inheriting maternal CD mutant but lacking any zygotic expression of MRG-1 (M-Z- WT; M+Z- CD) appear to lack any germ cell proliferation. Therefore unlike maternally supplied WT MRG-1, maternal CD mutant cannot rescue germ cell proliferation without further zygotic synthesis. The chromodomain is thus important for the early, maternally provided function of MRG-1, without which MRG-1 must be combined with early zygotic expression of either CD mutant or WT MRG-1 protein to at least partially support fertility. However, we found no obvious defects in post-embryonic germlines possessing only the CD mutant protein, suggesting the need for the chromodomain’s function in the germline is stage-specific

### Zygotic expression of *mrg-1* is necessary for normal somatic development

It has been shown that M-Z- *mrg-1(tm1227)* mutants lacking maternal MRG-1 fail to produce a germline due to defective proliferation of the PGCs(15, 17, 23). Here we describe a somatic role for MRG-1 in the sterile M-Z- *mrg-1* mutants. In addition to lacking a germline, a percentage of M-Z- mutants display severe somatic defects such stunted growth, tail defects, and multiple vulva development. This provides evidence that zygotically expressed MRG-1 plays an important role in somatic development. As a homolog of MRG15, that is known to influence developmentally regulated transcription, somatic defects in *mrg-1 M-Z-* mutant development are not surprising and are likely due to defective transcriptional regulation of developmental genes, presumably through its role(s) in histone modifying complexes. It is important to note, however, that despite the lack of either maternal or zygotic MRG-1, the *mrg-1(tm1227)* null allele survives embryogenesis and is thus either redundant with another factor or MRG-1 is not essential for early somatic development.

### The MRG-1 Chromodomain Mutations Cause Novel Phenotypes

The chromodomain of MRG-1 is highly conserved between species, so we characterized its role in *C. elegans* development by disrupting two of the five residues predicted to form a functional chromodomain aromatic “cage”. Interestingly, we found that the mutations caused a worse phenotype than the *tm1227* null allele: homozygous *mrg-1CD* mutants arrested and died as embryos. Since these homozygous CD/CD mutants arose from CD/WT mothers, the embryos inherited maternal WT MRG-1 and yet the maternal protein failed to rescue somatic development. Genetically, this would indicate that the CD mutant has a dominant, possibly antimorphic character that renders the maternal WT unable to rescue. However, since the null mutant shows that MRG-1 is not essential for embryonic viability, antagonism by the CD mutant of a WT protein that has no requirement in embryogenesis would seem to have few consequences (S7). Indeed, embryos that are of genotype *mrg-1^CD^/+ are* viable and exhibit no obvious meiotic defects, which indicates that presence of zygotically produced wild-type protein can suppress—or titrate away-- any negative effects of the CD mutant. This may be related to tissue specific functions of MRG-1 and the complexes in which it plays a role; the CD mutations may in effect induce a novel function to MRG-1 and its associated HDAC or HAT complex(es) important for embryonic gene regulation.

Surprisingly, the requirement for a functional CD is not observed in the post-embryonic germline. Using the AID system and auxin exposure starting in early larval stages, we were able to examine animals that had only the CD mutant present in their post-embryonic germline. These animals were fertile without any obvious germline defects and produced 100% dead embryos, as expected from the zygotic phenotype. Additionally, a majority of mrg-1^CD/WT::Degron^ embryos from *mrg*-1^CD/WT::Degron^ + auxin mothers that hatch on auxin (M-Z-WT; M+Z+ CD) grow up to be fertile suggesting that a combination of maternal and zygotic CD mutant MRG-1 can support fertility. This suggests the chromodomain of MRG-1is not essential for its role(s) in germline development.

Recently, MRG-1 has been identified a component of histone deacetylase complexes in *C. elegans* (12). Interestingly, mutations in the conserved histone deacetylase *hda-1* and a conserved complex member, *sin-*3, show embryonic lethality phenotypes similar to what we observed in the CD mutant embryos(30). The developmental defects caused by the CD mutations may be caused by changing how these complexes are targeted to genomic loci. Disruption of this targeting may alter deacetylase activity and its roles in regulating embryonic transcription.

### A Requirement for the MRG-1 Chromodomain in RNAi

We observed a strong RNAi defect in both CD/ + and CD/CD mutant animals, indicating that the RNAi defect is dominant. We do not know the direct cause of the RNAi defect, but propose that it is likely an indirect effect. As mentioned above, MRG-1 has been identified by mass-spectrometry as a component of numerous chromatin-modifying complexes in *C. elegans* (12, 13).The defective chromodomain could affect any role MRG-1 may have in the targeting of these complexes to their genomic substrates, which in turn could result in defective transcriptional regulation of RNAi machinery. Alternatively, defective histone acetylation dynamics could lead to defective suppression of endogenous RNAi targets, thereby causing endogenous and exogenous RNAi systems to compete for shared factors in their pathways. These phenotypes could be the result of altered acetylation and transcriptional patterns that are regulated by HDACs and/or HATs through MRG-1’s chromodomain, or dominantly affected by any novel activity induced by the CD mutations. Indeed, mutations in HDAC complexes have been shown to have defective RNAi phenotypes, similar to the chromodomain mutant(31). It is also interesting to note that *mes-*4 M+Z- animals were shown to have an RNAi defect, which may point to H3K36 methylation, and possibly its recognition by MRG-1 via its chromodomain, as important for normal expression of RNAi effectors(32, 33). Importantly, whatever role the chromodomain has in transcription regulation is not evident in the post-embryonic germline, either because it is not required or is redundant with other histone modification readers. Further analyses of the CD mutant and its effects are obviously warranted and are in progress.

## Materials and Methods

### Strains and Strain Maintenance

The following strains were used for these experiments: Bristol N2, XA6227 (*<u>mrg-1</u>*[*<u>tm1227</u>*]/qC1 [*<u>dpy-19</u>*(*<u>e1259</u>*) *<u>glp-1</u>*(*<u>q339</u>*) qIs26] III), AV630 (<u>meIs8</u> [pie-1p::GFP::cosa-1 + unc-119(+)] II), MT16973(*met-1*[<u>n4337</u>] I). Additional transgenic strains were generated using the CRISPR-Cas9 system described below. All strains were maintained at 20°C on nematode growth medium (NGM) plates seeded with *E. coli* OP50. For *mrg-1(tm1227)* mutants, all analyses were performed in the M+Z- (maternal +/ zygotic -) generation. Some strains were provided by the CGC, which is funded by NIH Office of Research Infrastructure Programs (P40 OD010440). Any newly generated strains will be made available through the CGC.

### CRISPR Cas9 Generated Strains

The endogenous *mrg-1* gene was tagged C-terminally tagged with mCherry, GFP, or AIDdegron::GFP using the CRISPR-Cas9 genome editing system (described below)(26). Specific guide RNAs were used to induce breaks in the last exon of *mrg-1* and were repaired using short DNA gene fragments (Integrated DNA Technologies) containing the specific protein label. The CRISPR-Cas9 system was also used to mutate chromodomain of *mrg-1* in a similar manner with specific guide RNAs (gRNAs) and a repair oligonucleotide (S Table 2-3). Large gBlock gene fragments were amplified using Platinum SuperFi PCR Master Mix (Invitrogen #12358010) and concentrated using the Zymo DNA Clean & Concetrator-5 kit (Zymo D4004) before injection young adult worms.

| Strain | Description |
| --- | --- |
| KW1668 | ck42(*mrg-1::mcherry*) III |
| KW1681 | ck43:(*mrg-1::mcherry* CD mutation); mutated Y17A & F43A/<br>qC1[dpy-19(e1259) glp-1(q339) qls26] III |
| KW1686 | ck44(*mrg-1::GFP*) III |
| KW1682 | ck45(*mrg-1::degron::GFP*) III;<br>ieSi38 [sun-1p::TIR1::mRuby::sun-1 3'UTR + Cbr-unc-119(+)] IV |
| KW1683 | *mrg-1(ck43)/mrg-1(ck45)* III; ieSi38 IV |

### RNAi Conditions

RNAi was performed by using HT115 cells transformed with the empty L4440 RNAi vector or L4440 vector containing the cDNA of the gene of interest. Bacteria was grown shaking overnight at 37°C. Bacteria cultures were induced with IPTG 1mM final concentration for 1 hour while shaking and plated on NGM plates with ampicillin (100ug/mL) and IPTG(1mM). Bacteria was induced on ampicillin-IPTG plates overnight at 37°C degrees. After at least 36 hours of induction, synchronized L1s were plated on the RNAi bacteria plates and then moved to 20°C until analyses.

### Synchronization

To synchronize worm populations, worms were grown on NGM plates until gravid adults. Animals were rinsed off the plates with M9 buffer, collected in a 15ml conical tube, and spun down at 1800rpm for 3 min. Once pelleted, M9 was discarded leaving 6X of worm pellet volume (up to 3mL). The pellet was resuspended with 10% volume of 50% bleach solution and 10 N NaOH and incubated on a shaker until worms were dissolved. The released embryos were then washed with 15ml M9 three times. After the last wash, the embryo pellet was resuspended in M9 buffer and placed on a shaker overnight for the embryos to hatch. The synchronized L1 larvae were then plated on seeded NGM plates.

### Immunofluorescence

Germlines were dissected from young adult (24 hours after L4 larval stage) hermaphrodite worms in dissection buffer (2X sperm salts, 2.5mm levamisole) on poly-lysine coated slides and fixed with 1% paraformaldehyde with ethanol or methanol acetone following freeze crack on a chilled aluminum block. After fixation, slides were washed in TBST (tris buffer saline, 1% Tween-20) and primary antibody was added for incubation at 4° overnight. Following washes with TBST, slides were incubated with secondary antibody at room temperature for 4 hours. Slides were then washed with TBST, and then counterstained with DAPI. Slides were sealed with Prolong Gold and clear nail polish. Imaging was done using a Leica DMRXA with a Retiga2000r camera using Hamamatsu Photonics software. Confocal images were taken with an Olympus FV1000 confocal microscope using Olympus Fluoview v4.2 acquisition software. This research project was supported in part by the Emory University Integrated Cellular Imaging Core. The content is solely the responsibility of the authors and does not necessarily reflect the official views of the National Institute of Health. All image analysis was done using FIJI imaging software(34).

The following primary antibodies were used: goat anti-SYP-1 (1:1500), chicken anti-GFP (Aves Labs GFP-1020, 1:300), mouse anti H3K36me3 (Active Motif 61020, 1:20000), Secondary antibodies used in this study were: donkey anti-goat 488 (Invitrogen A11055, 1:500), donkey anti-mouse 594 (Invitrogen A21203, 1:500), donkey anti-chicken 594 (Jackson Immuno Research Laboratories 703586155, 1:500).

### Brood Size Assay

Individual L4 larvae were placed on NGM plates seeded with OP50. After 24 hours, worms were transferred to new plates. Embryos and L1s were then counted and the number of unhatched embryos were scored the following day to determine embryonic lethality. This was repeated until embryos were no longer produced. Brood size was calculated as the total number embryos laid. Embryonic lethality was determined by the percentage of embryos in each brood that failed to hatch after 24 hours.

### Transition Zone Measurement

Transition zone (TZ) nuclei were identified in dissected, fixed and DAPI stained gonads as nuclei with characteristic crescent shaped and condensed chromosomes asymmetrically located to one side of the nuclear periphery. The length of the TZ was measured by counting the number of linear nuclei from the first distal crescent-shaped DAPI nucleus to the most proximal. TZ length analysis was done by comparing the nuclear length of the TZ to the total nuclear lengths from the first nucleus in the distal mitotic region to the last nucleus at the end of pachytene before diplotene. The Mann-Whitney U-test was used for statistical analysis (p =2.677E-05).

### Crossover Formation Characterization

After RNAi, dissected gonads from AV630 were stained for GFP and SYP-1. Maximum projections of Z-stacks of each gonad were analyzed for crossover formations. The number of crossovers formed in each nuclei were determined by the number of GFP foci in each nuclei. GFP::COSA-1 foci were only counted if they colocalized with DAPI staining. The length of the gonad was divided into six sections and the number of GFP::COSA-1 foci per nucleus were noted in each section. Significance was calculated using the Chi-Square test comparing the type of nuclei present in region (Region 5: p= 2.03E-04, Region 6 p= 2.01E-05).

For bivalent and univalent formation analysis, the number of DAPI bodies were counted in oocytes between wildtype (n=121) and *mrg-1 M+Z-* (n=61). The Chi Square test was used for statistical analysis between control and *mrg-1*(p = 3.40275E-14).

### Auxin Induced Degradation

Auxin Induced Degradation (AID) experiments were performed by the method of (26). OP50 seeded NGM plates were supplemented with 1mM of auxin and the bacteria lawn was allowed to grow for at least 48 hours at room temperature before worms were added to plates. Worms were placed on auxin containing plates for the indicated amount of time before being analyzed further.

## Acknowledgements

We would like to thank S. L’Hernault for CRISPR design suggestions. We would like to thank all of the past and current members of the Kelly Lab and Deal Lab for their helpful discussion and comments during the scope of this project.

## Supplemental Figures

**S1.**
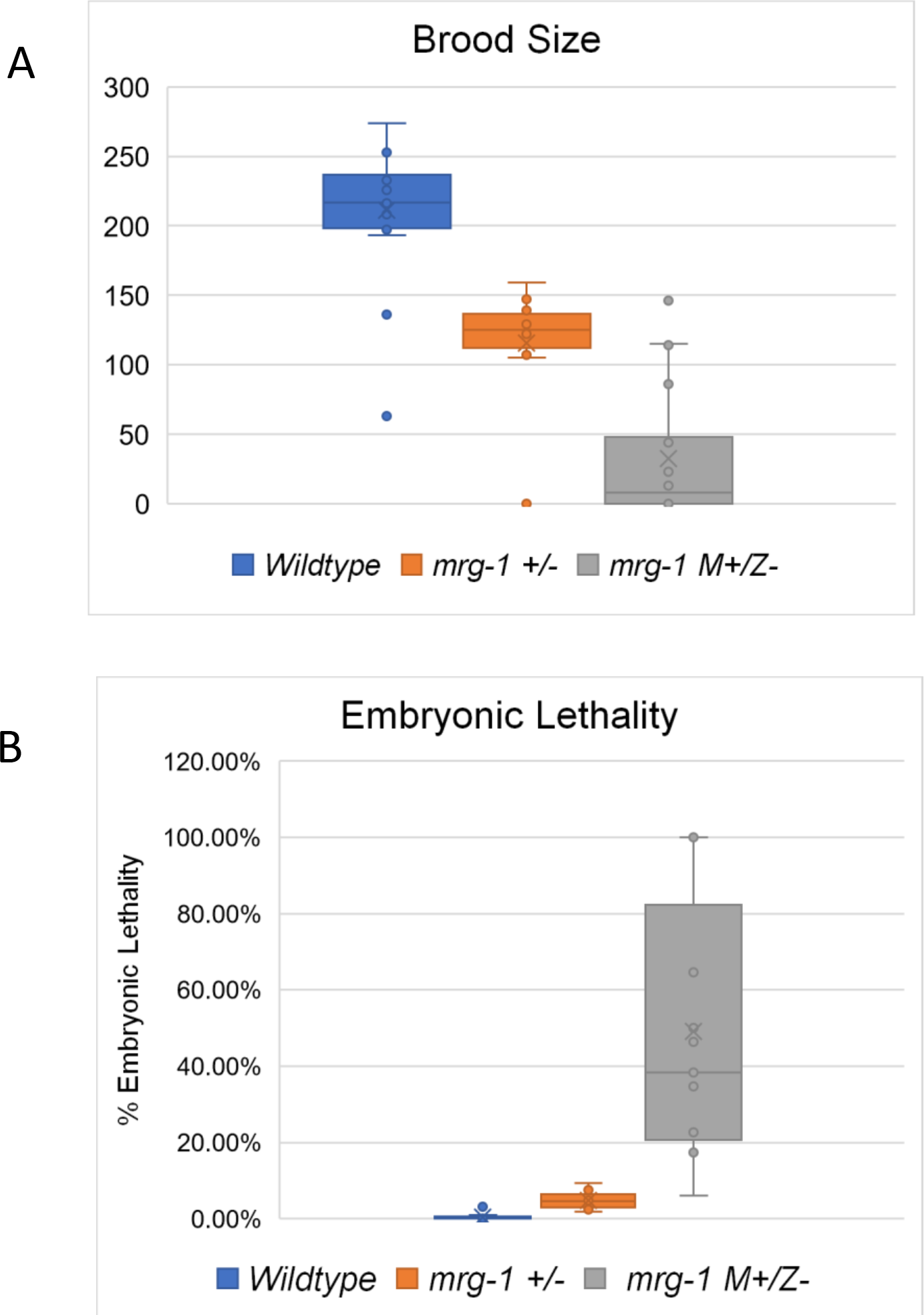
*mrg-1 (tm1227)* mutants have decreased brood size and increased embryonic lethality. The total brood of wildtype, *mrg-1 (tm12227)* heterozygotes, and *mrg-1 M+Z-* worms were counted (A), and the percentage embryos failing to hatch was calculated (B). *mrg-1 M+Z-* worms had a higher percentage of embryonic lethality compared to wildtype and *mrg-1* heterozygotes. The broods of 25 animals were counted for each genotype.

**S2.**
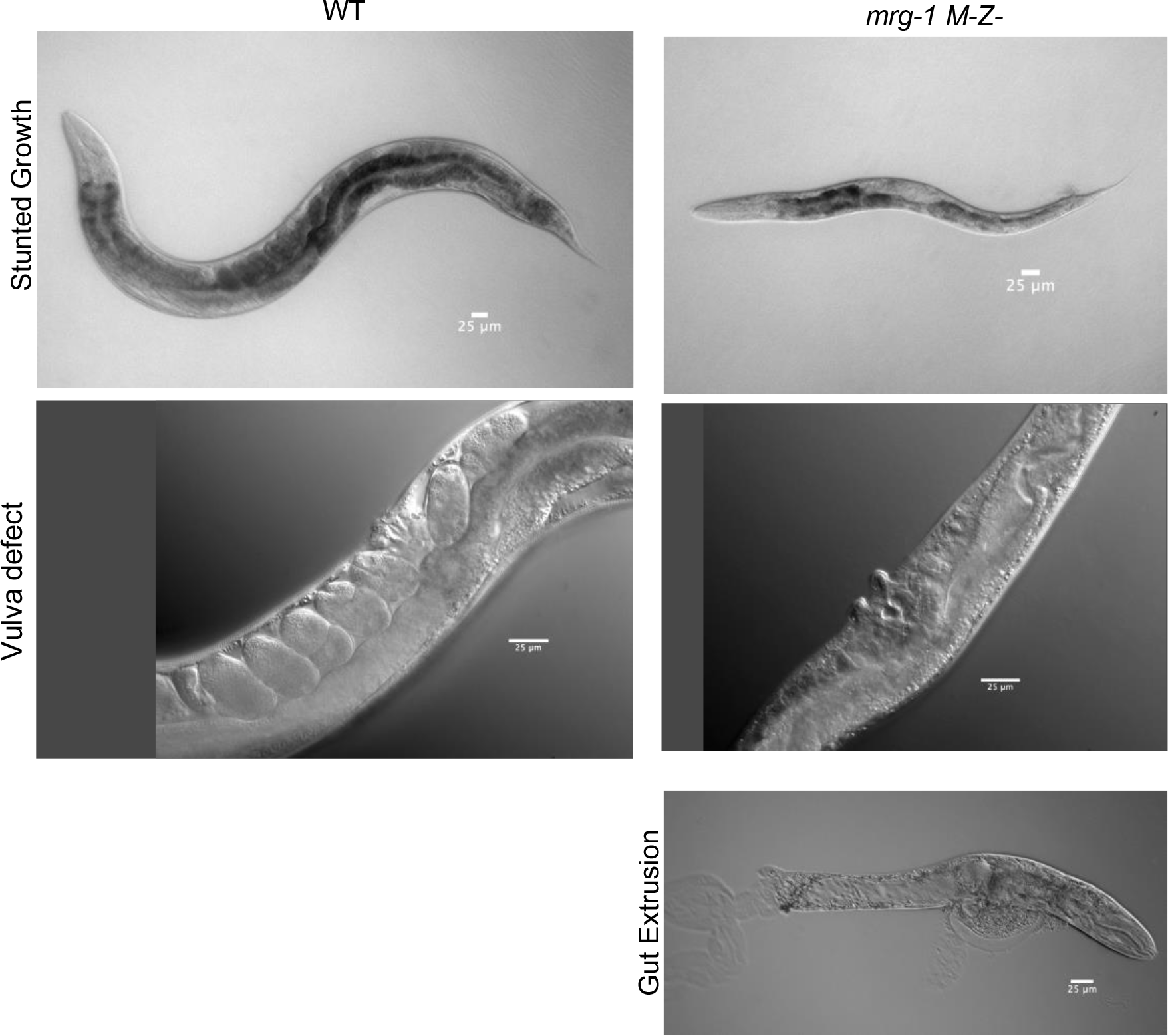
*mrg-1 M-Z-* mutants exhibit somatic defects. M-Z- progeny of M+Z- *mrg- 1(tm1227)* mutants were imaged 24 hrs after the L4 stage. A majority of the worms displayed a variety of somatic defects including stunted growth, protruding vulvae, and extrusion of the gut through the anus. The worms that did not display somatic defects were sterile as expected.

**S3.**
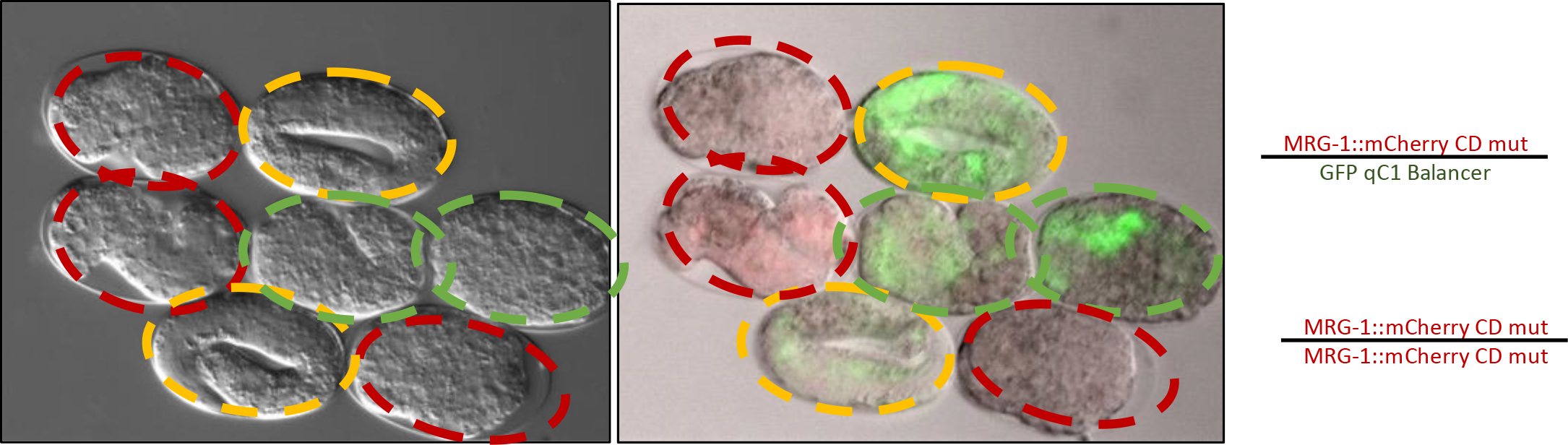
MRG-1 Chromodomain mutation causes embryonic arrest. Embryos dissected from *mrg-1CD/qC1* balancer worms were dissected and allowed to develop. Embryos that were homozygous for *mrg-1CD::mCherrry* mutation lack GFP and arrested during embryogenesis (red outlines). Heterozygous siblings (express GFP and mCherry) complete embryogenesis, hatch and develop normally (yellow outlines). Embryos that are homozygous for the qC1 balancer also arrest (green outline)

**S4.**
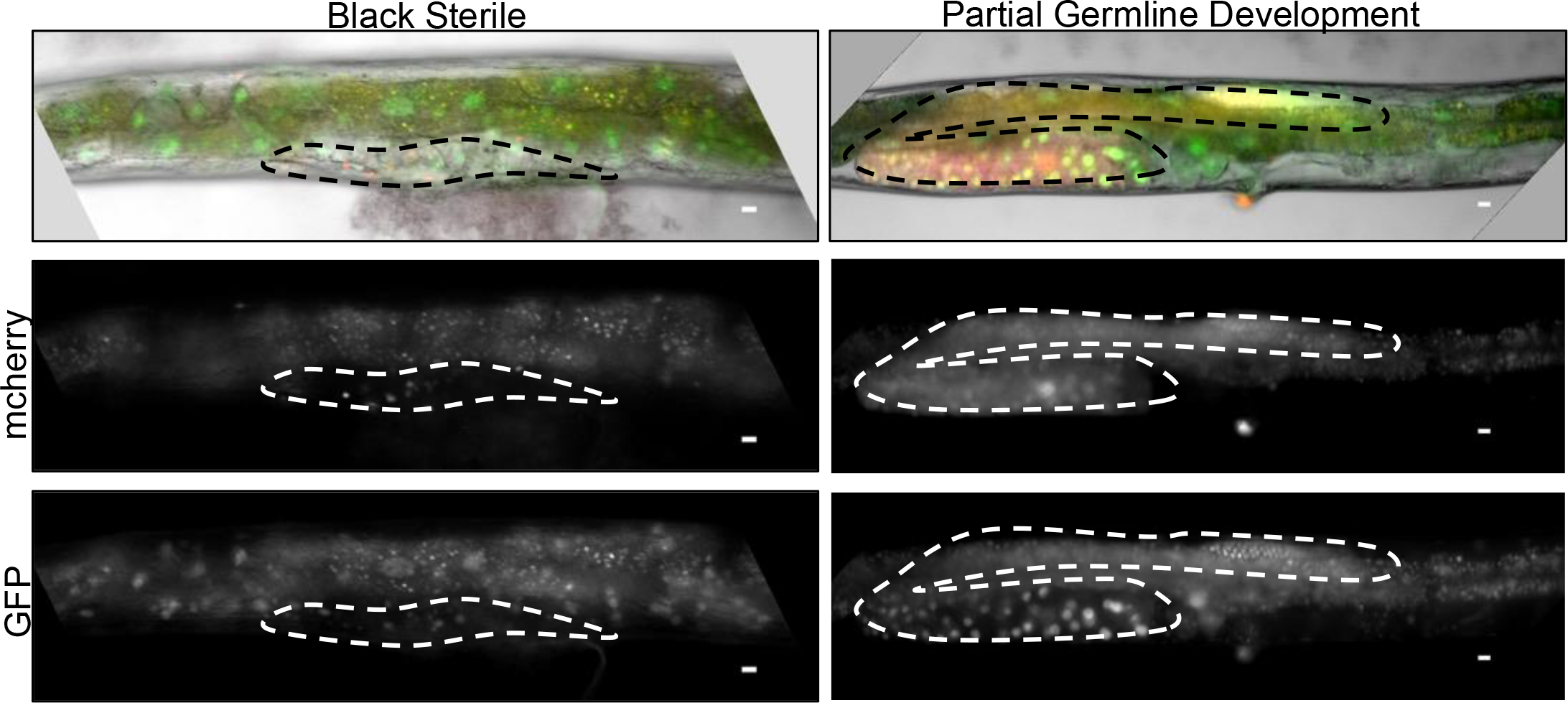
Zygotic expression of WT MRG-1 partially rescues germline development. Sterile worms that had zygotic expression of WT or WT and CD mutant, had various levels of sterility. Some worms completely lacked germ cell proliferation and were black sterile (left), while other worms had expression of both WT and CD with germline development but still lacked fertility (Right).

**S5.**
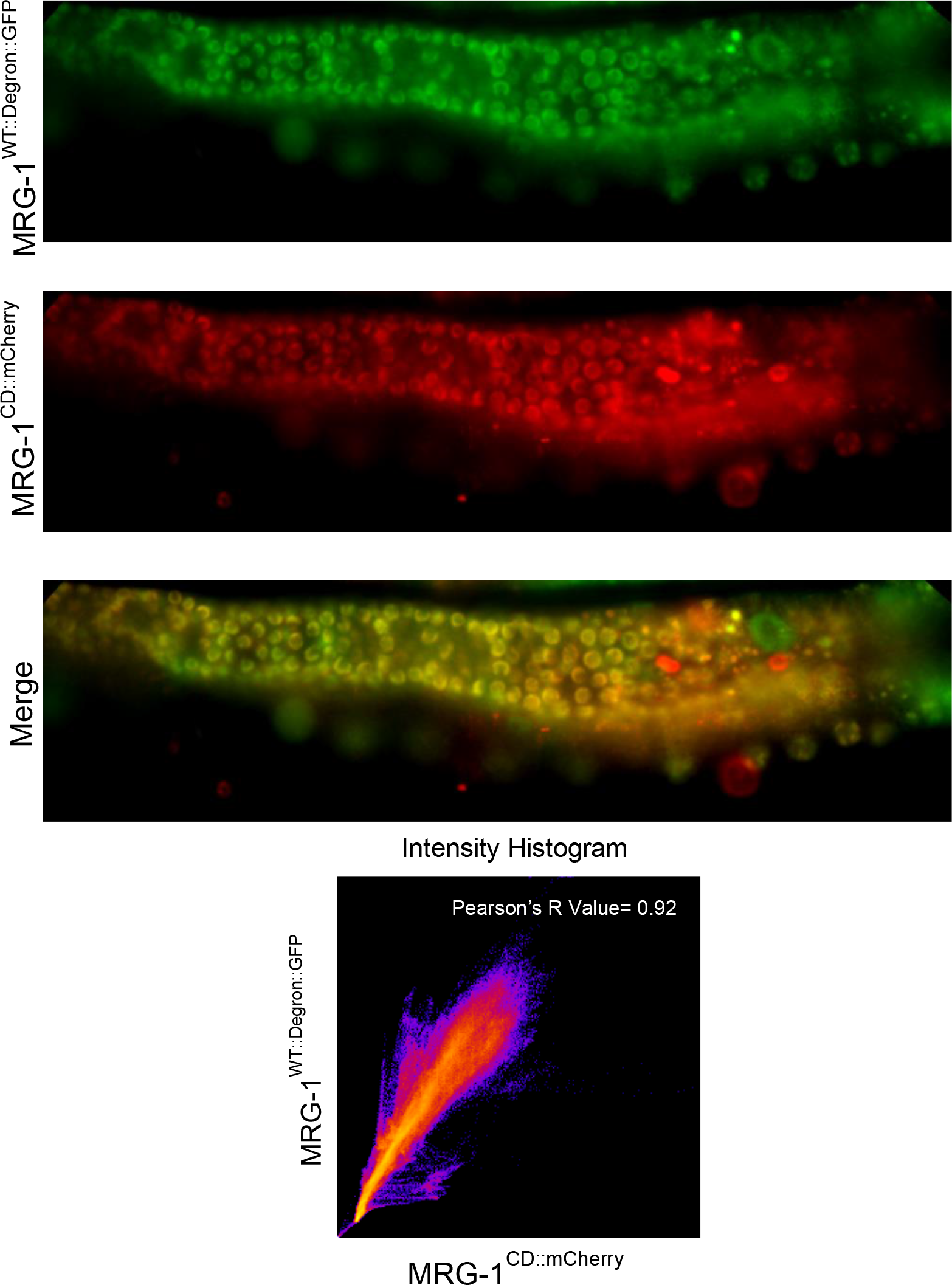
MRG-1^CD::mcherry^ colocalizes with MRG-1^WT::Degron::GFP^. Live images of *mrg- 1^CD/WT::Degron::GFP^* worms were taken. MRG-1^CD:;mCherry^ has the same localization pattern as MRG-1^WT::Degron::GFP^ in the germline. Colocalization analysis was performed and resulted with a Pearson’s R value of 0.92.

**S6.**
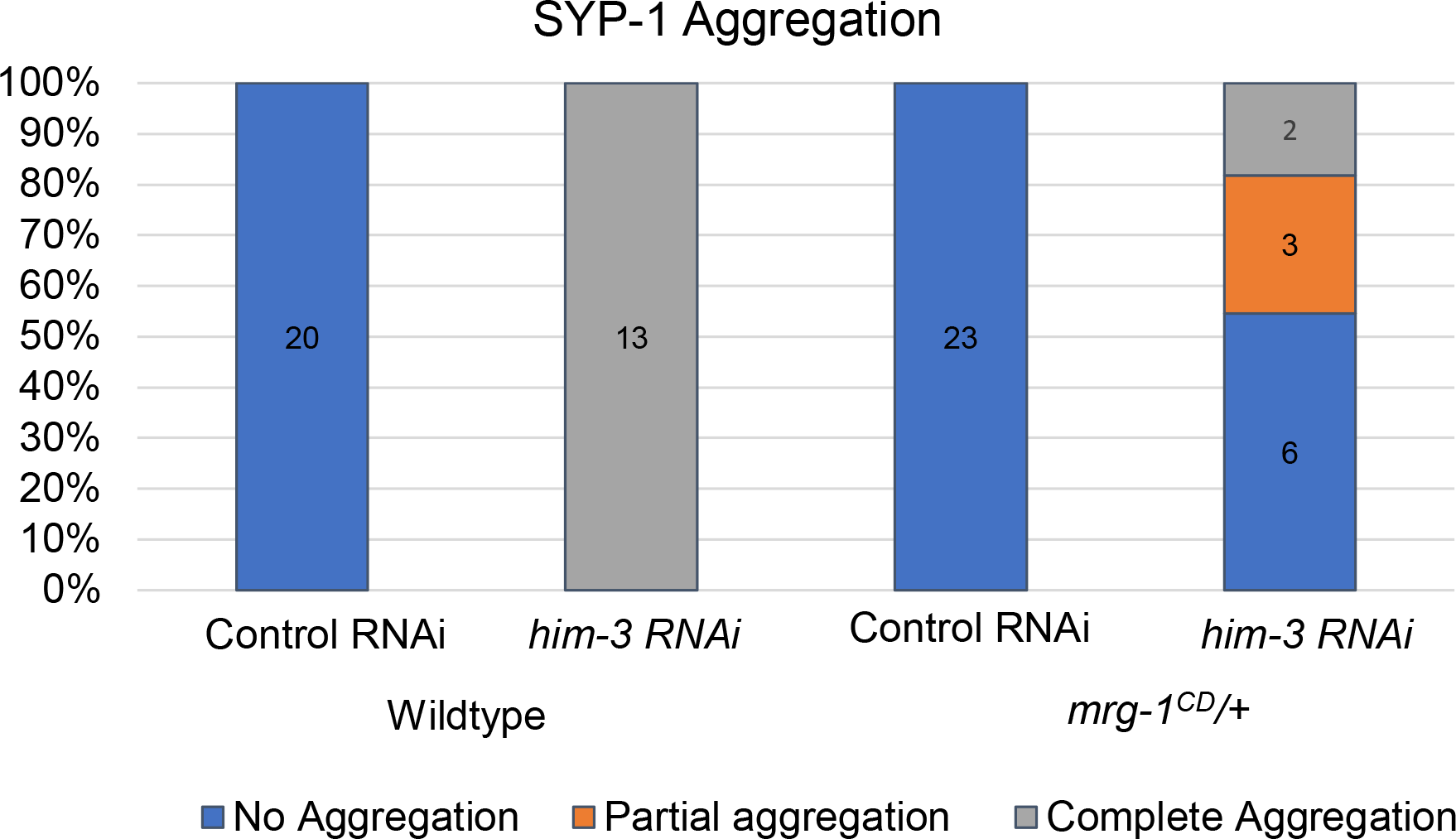
*mrg-1 CD/+* hermaphrodites exhibit some SYP-1 aggregation after *him-3 RNAi.* WT and *mrg-1 CD/+* hermaphrodites were placed on control or *him-3* RNAi from L1 Larval stage. SYP-1 aggregation was seen all WT germlines after *him-3* RNAi. In *mrg- 1CD/+* germlines, a range of aggregation phenotypes were seen from zero to complete aggregation.

**S7.**
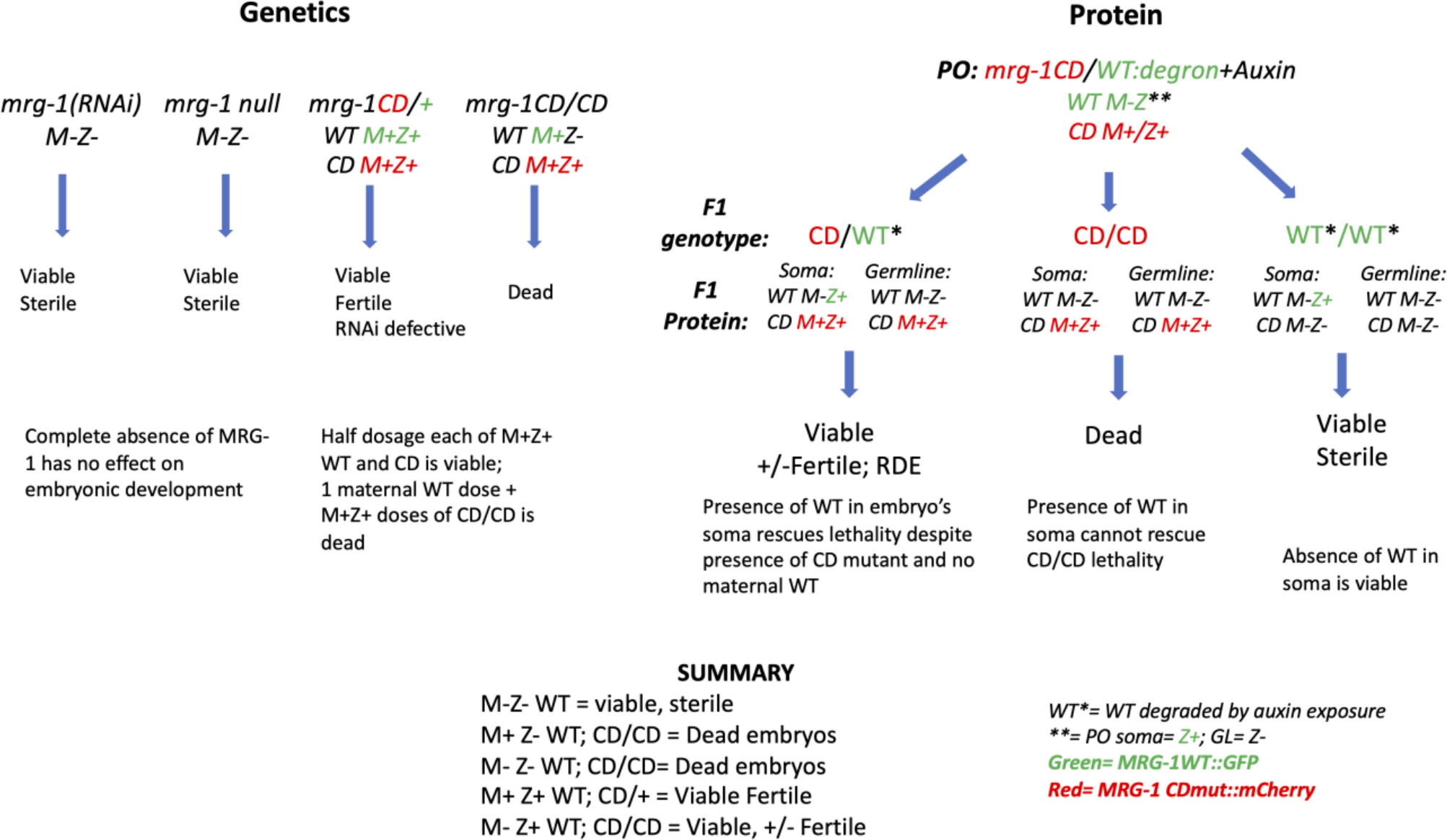
Summary of Genetic and Protein Phenotypes of the *mrg-1(tm1227)* and *mrg- 1^CD^* Mutant. The phenotypes observed from the *tm1227* allele and those observed in the AID experiments with degron-tagged WT MRG-1. The presence or absence of maternally provided (M) or zygotically produced (Z) MRG-1 is indicated. The presence of GFP-tagged WT MRG-1 protein (Green) and/or mCherry tagged CD mutant MRG-1 protein (Red) are indicated. The RNAi results are from ref. 14.

**S Table 1.** *mrg-1* M-Z- Somatic Defects.

| ***mrg-1* M-Z- Somatic Defects** |  |
| --- | --- |
| **Phenotype** | **Percentage (n = 51)** |
| Stunted Growth | 21.6 % (11) |
| Protruding Vulva | 51.0% (26) |
| Gut Extrusion | 13.7%(7) |
| WT | 13.7 % (7) |

**S Table 2.**
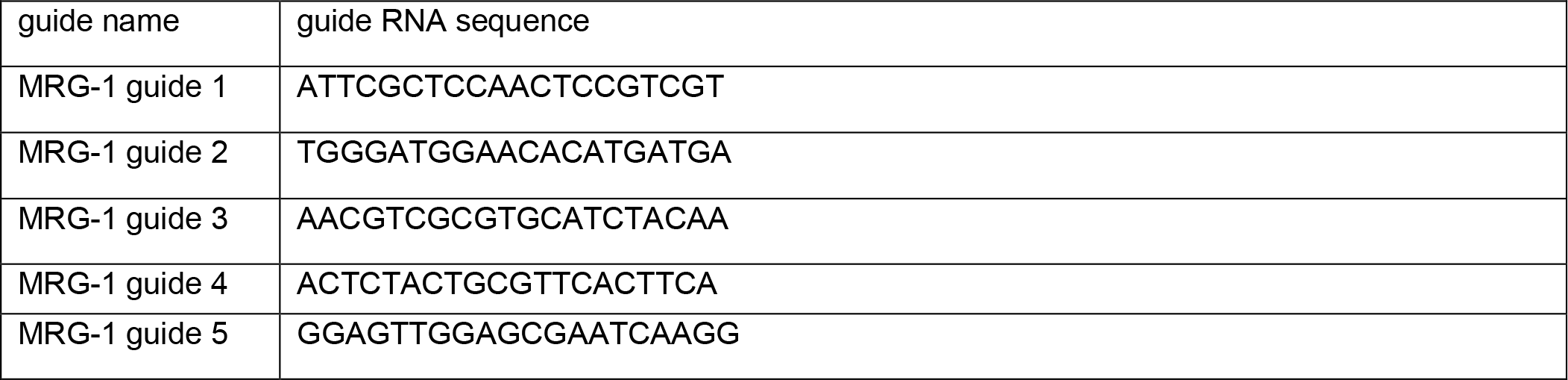
Guide RNA Sequences. Guide RNA sequences used to induce breaks at the endogenous *mrg-1* locus using the CRISPR-Cas9 system.

**S Table 3.**
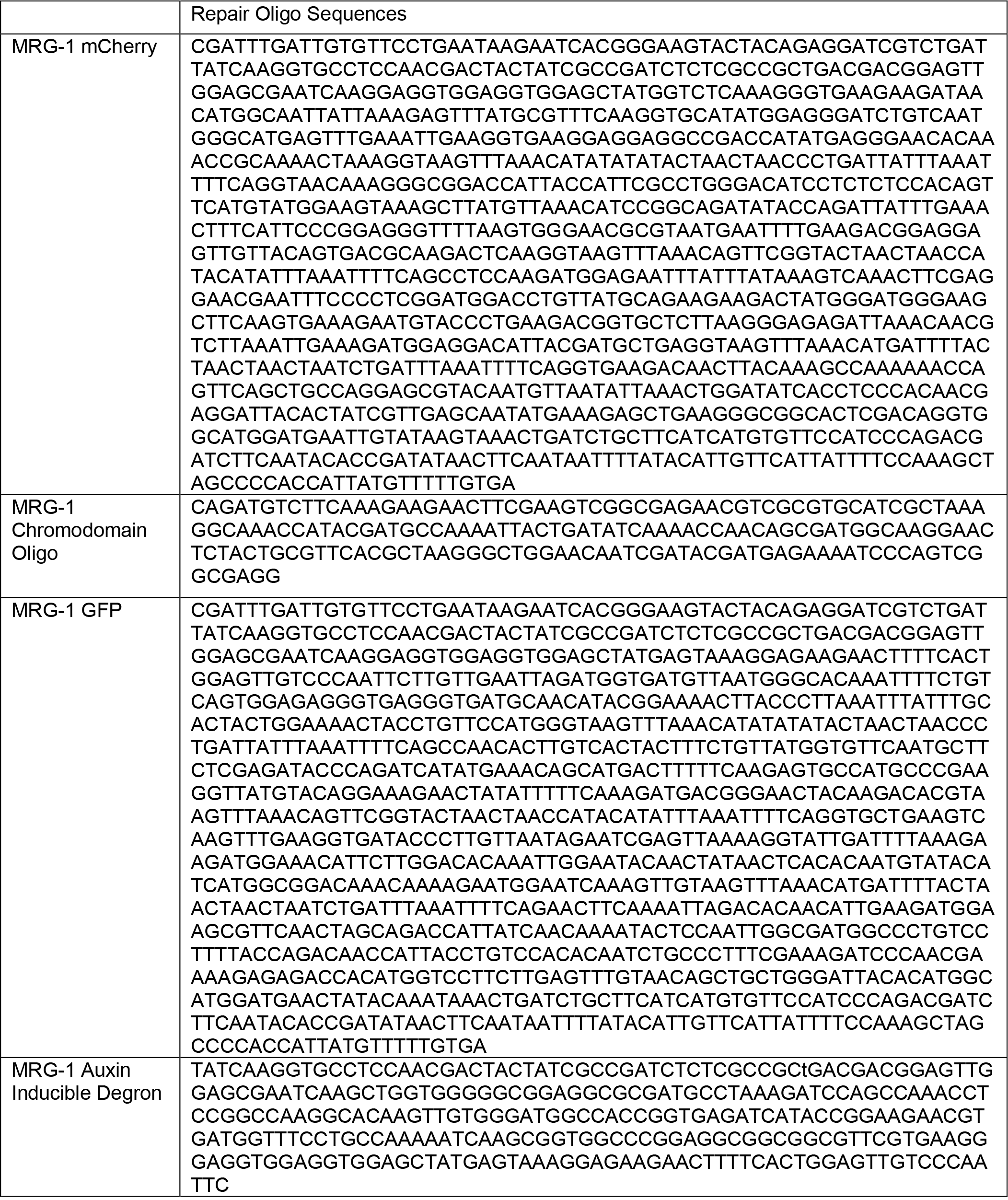
Oligonucleotide Repair Sequences. Oligonucleotide sequences used as repair sequences to generate specified transgenic strains using the CRISPR-Cas9 System.

